# Integrating diverse data sources to predict disease risk in dairy cattle

**DOI:** 10.1101/2021.03.25.436798

**Authors:** Jana Lasser, Caspar Matzhold, Christa Egger-Danner, Birgit Fuerst-Waltl, Franz Steininger, Thomas Wittek, Peter Klimek

## Abstract

Livestock farming is currently undergoing a digital revolution and becoming increasingly data-driven. Yet, such data often reside in disconnected silos making it impossible to leverage their full potential to improve animal well-being. Here, we introduce a precision medicine approach, bringing together information streams from a variety of life domains of dairy cattle to predict eight common and economically important diseases. Dairy cows are part of a highly industrialised environment. The animals and their surroundings are closely monitored and environmental, behavioural and physiological observations are readily accessible yet seldomly integrated. We use random forest classifiers trained on data from 5,828 animals in 166 herds in Austria to predict occurrences of lameness, acute and chronic mastitis, anoestrus, ovarian cysts, metritis, ketosis (hyperketonemia) and periparturient hypocalcemia (milk fever). To assess the importance of specific cattle life domains and individual features for these predictions, we use multivariate logistic regression and feature permutation approaches. We show that disease in dairy cattle is a product of the complex interplay between a multitude of life domains such as housing, nutrition or climate, and identify a range of features that were previously not associated with increased disease risk. For example, we can predict anoestrus with high sensitivity and specificity (F1=0.72) and find that housing, feed and husbandry variables such as barn design and time on pasture are most predictive of this disease. We also find previously unknown associations of features with disease risk, for example humid conditions, which significantly decrease the odds for ketosis. Our findings pave the way towards data-driven point-of-care interventions and demonstrate the added value of integrating all available data in the dairy industry to improve animal well-being and reduce disease risk.

## 1 Introduction

During the previous decades, precision medicine for humans has been recognised as one of the most promising new approaches to understand health and disease^1–3^. In precision medicine, information from many sources is combined to get a holistic picture of an organism and its surroundings and find tailor-made treatments for diseases. In livestock farming, precision medicine has been conducted under the umbrella of precision livestock farming^4, 5^ (PLF). As the health of livestock has large economic implications, the determination of risk factors for diseases is a frequent application in PLF. Additionally, PLF holds great promise to steer livestock farming into a more environmentally sustainable direction^5, 6^ by enabling preventive interventions and reducing animal losses. Animal well-being is increasingly being considered an important economic factor as well, as consumers get more conscious about the origins of the products they buy^7, 8^. Since dairy cows are part of a highly industrialised environment, the animals and their surroundings are closely monitored and environmental, behavioural and physiological observations are measured in routine assessment^9, 10^. The digitisation and integration of information from these different sources has large potential to determine risk factors and arrive at actionable information allowing. Therefore, the main goal of this study is the improved health and herd management on dairy farms by enabling individualised data-driven point-of-care interventions through the integration of information from a multitude of different sources around a farm.

Previous studies have often focused on finding relationships between isolated areas of dairy cow husbandry, such as nutrition or milk parameters and disease outcomes (see for example Oehm et al. 2019^11^, Roche et al. 2019^12^ and Polsky et al. 2017^13^ for some recent reviews). In this publication, we show that aggregating data from a range of diverse sources to predict diseases in dairy cattle improves prediction accuracy compared to predictions using less diverse data, leads to new insights about important factors that influence disease incidence and creates value by re-using existing data pools^14^. We show that no single factor predicts disease incidence alone. Diseases are indeed a result of a complex interaction between multiple influencing factors from all life domains of the animals.

Several machine learning (ML) approaches have been considered for the prediction of breeding values^15^, insemination success^16^, feed intake^17^ and calving^18^, as well as milk yield^19–21^, modelling of physiological and behavioural animal parameters^22, 23^ and estimating BCS^24, 25^ – see also Cockburn 2020^26^ for a recent review. The literature on the prediction of diseases is frequently based on black-box sensor systems in which prediction algorithms are used that are of commercial interest and not publicly known or evaluated^26^ or the reported sensitivity and specificity of prediction algorithms varies widely^9^. Diseases for which successful applications of ML approaches have been reported include lameness^27–29^, mastitis^30–33^, metabolic status, i.e. ketosis (hyperketonemia) and periparturient hypocalcemia (milk fever)^34–38^, infectious diseases^39, 40^ and oestrus detection^41^. To our knowledge, no studies describing the application of ML approaches for the prediction of anoestrus, metritis and ovarian cysts exist to date.

We analyse 22,923 observations of 5,828 animals on 166 dairy farms in Austria from the years 2014 to 2016. Each observation has 138 features, derived from eleven different life domains of the animal, showcased in fig. 1. Our features include information that has a long been known as relevant for the assessment of dairy cattle fitness and disease prediction, such as breeding values and information about animal breeds^42^ and dairy herd improvement (DHI) assessment information expanded by body and conformation assessments^43, 44^. We combine this information with extensive information on husbandry and management conditions of the cattle, such as barn architecture and manure removal practices, as well as farm management, such as type and duration of pasturing and information about the milking system. Furthermore, we add detailed information about feed composition, such as the dietary proportion of crude fibre and concentrates and metabolic indicators. We use information from weather services to add environmental factors such as the average rainfall and temperature. Lastly, we add information about the current lactation state of the animal as well as its parity.

**Figure 1.**
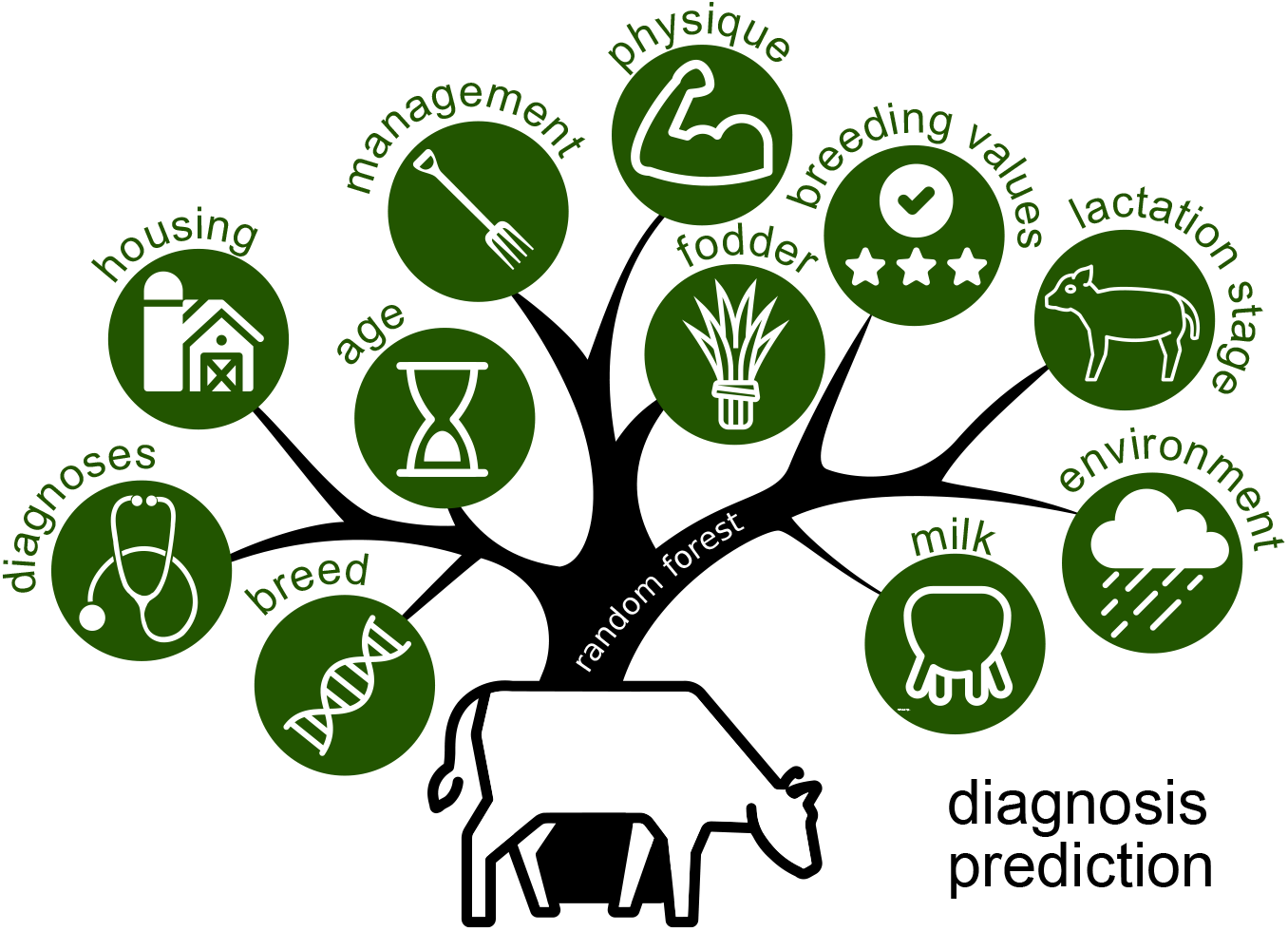
Eleven sources of information, shown here as leafs on a tree, are aggregated to predict common dairy cattle diseases using a random forest classifier^45^.

This information is used to predict a range of common dairy cattle diseases that are of high economic relevance, such as ketosis, lameness and mastitis. Information about the occurrence of these diseases are obtained from diagnoses by veterinarians, observations at calving or culling reasons. In addition, we add observations based on lameness scoring and ketosis tests to label diseases (see sec. 4.1). We will refer to all these disease labels as “diagnosis” in the remainder of this work, regardless of origin.

We train a random forest classifier on these diagnosis labels to predict the corresponding disease in unlabelled data, i.e. in data in which we removed the diagnosis information and which the algorithm has not seen during training. To train the random forests, we use a total of 138 features from 11 categories aggregated from 6 different sources (see supplement table 1 for an overview of the available features and data origin). The training routine for the random forests are described in section 4.2. Furthermore, for all diagnoses we test two versions of our data set: one including all observations, and one including only observations from animals which were lactating at the time of observation (19703 observations). The reasoning behind this division is the large metabolic change a cow undergoes between lactation and dry periods. Therefore observations from the dry period might not be suited to predict diseases that occur during lactation.

## 2 Results & discussion

In the following, we report our results on the quality of the predictions of different diseases and discuss how these findings relate to other machine learning approaches proposed in the literature. We then report which variables and life domains showed a particularly high predictive value for different diseases and discuss these results.

### 2.1 Disease prediction accuracies

Our data set contains 22,923 observations of healthy (59.6% of observations) and sick cows (40.4% of observations), including information on herd living and environmental conditions^46^, individual cow feed^47^, milk and physical parameters such as feed energy content, test-day milk yield or animal weight, and 56 diagnosis codes (see sec. 4.1 and^46^ for more details on the data). The eight most prevalent diagnoses in our data set are lameness (39.6% of all diagnoses), acute mastitis (11.2%), anoestrus (8.7%), ovarian cysts (7.6%), periparturient hypocalcemia (7.3%), ketosis (7.1%), chronic mastitis (3.6%) and metritis (3.4%; see tab. 1 for diagnosis frequencies). Diagnoses frequencies in our data set do not reflect true prevalences of the respective diseases in dairy cattle. This is due to the aggregation of diagnoses data with several other data sets of observations through matching of observation and diagnoses dates (see sec. 4.1) During the matching process, diagnoses that do not have matching observations are excluded. Data on true disease prevalences in a comparable setting is given in Egger-Danner et al. 2012^48^. Lameness is most likely over-represented in our data, since we combine the clinical diagnoses with lameness scores to create a positive diagnosis label. Similarly, we combine clinical ketosis diagnoses with subclinical diagnoses based on elevated ketone body levels in ketosis tests. Nevertheless, since animals subjected to ketosis tests are not drawn from a representative sample, the number of subclinical ketosis diagnoses is not representative and the number of diagnoses is on the lower end of the prevalence spectrum reported in literature, which ranges from 6.6 %^49^ to 47.2 %^50^.

**Table 1.** Frequency of diagnoses for the eight most frequent diseases. The percentages denote the share of the diagnoses among all observations with a diagnosis contained in the data set.

| diagnosis | N observations | % |
| --- | --- | --- |
| lameness | 3670 | 39.6 |
| acute mastitis | 1037 | 11.2 |
| anoestrus | 803 | 8.7 |
| ovarian cysts | 705 | 7.6 |
| periparturient hypocalcemia | 673 | 7.3 |
| ketosis | 657 | 7.1 |
| chronic mastitis | 335 | 3.6 |
| metritis | 318 | 3.4 |
| other | 1062 | 11.5 |

Prediction sensitivity, specificity and F1 score for the eight selected diseases and two data set versions (dry and lactating cows or only lactating cows) are listed in Tab. 2. All our classifiers generally achieve high prediction specificities due to the low prevalence of most diagnoses and the high number of true negatives. Of higher interest are the sensitivities, which are typically substantially lower than the specificities. Prediction performance differs between the two data set versions. Predictions achieve the highest F1 score for anoestrus in both versions (F1 score of 0.720 excluding dry period and 0.731 in the full data set, respectively). Similar prediction accuracies for anoestrus in both data sets are expected, since anoestrus cannot occur in dry cows. As the only existing other study^51^ that investigates anoestrus in dairy cattle focuses on the the detection of oestrus behaviour rather than the prediction of anoestrus, a direct comparison between prediction performance is impossible. It is worth noting that the predictor achieves a similarly high F1 score for anoestrus as for lameness, even though the amount of available observations is less than a third of the observations available for lameness.

**Table 2.** Prediction F1 scores, sensitivity, specificity, diagnosis prevalence and number of observations for the eight most prevalent diseases in our data set. The random forests were trained with hyper-parameters optimised for each disease group (see supplement table 2).

| diagnosis | excluding dry period |  |  | full data set |  |  | N diagnoses |
| --- | --- | --- | --- | --- | --- | --- | --- |
|  | F1 score | sensitivity | specificity | F1 score | sensitivity | specificity |  |
| anoestrus | 0.720 | 0.635 | 0.986 | 0.731 | 0.635 | 0.989 | 753 |
| lameness | 0.672 | 0.564 | 0.951 | 0.715 | 0.631 | 0.939 | 3361 |
| ketosis | 0.449 | 0.317 | 0.994 | 0.701 | 0.678 | 0.975 | 355 |
| acute mastitis | 0.327 | 0.203 | 0.995 | 0.416 | 0.277 | 0.992 | 830 |
| chronic mastitis | 0.452 | 0.312 | 0.996 | 0.339 | 0.291 | 0.981 | 312 |
| ovarian cysts | 0.464 | 0.411 | 0.966 | 0.482 | 0.433 | 0.961 | 668 |
| periparturient hypocalcemia | 0.275 | 0.175 | 0.995 | 0.619 | 0.553 | 0.978 | 293 |
| metritis | 0.500 | 0.490 | 0.980 | 0.611 | 0.496 | 0.994 | 265 |

Prediction of lameness works similarly well in both data sets (F1 score of 0.672 vs. 0.715) with a sensitivity of 0.631 in the full data set. Sensitivity is similar to Ghotoorlar et al.^27^ and slightly outperforms the approach presented by Warner et al.^28^. Deep-learning based lameness detection^29^ with videos of moving animals as input clearly outperform all other approaches with a sensitivity of 97.51%.

Prediction of acute and chronic mastitis only achieves moderate F1 scores due to low sensitivity in both data sets (F1=0.416 and F1=0.339 in the full data set and F1=0.327 and F1=0.452 in the reduced data set which excludes lactating cows), even though the number of observations with these diagnoses is comparable to the number of observations of anoestrus. Combining both diagnoses into a general “mastitis” diagnosis did not improve prediction sensitivities. It is worth noting that chronic mastitis is the only condition for which the predictor trained on the reduced data set yielded a higher sensitivity than the one trained on the full data set. Two of the four existing studies on the application of ML algorithms to mastitis phenomenology^30, 31^ do not predict disease incidence and a direct comparison with our approach is not possible. The other two approaches outperform our approach with sensitivities of >93%^33^ and 97.7%^32^ respectively. Both studies use neural net based approaches applied to milking information. Information about milking systems and routines – which is present in our data set – does not seem to appropriately capture the causes for mastitis. We assume that adding information about hygiene and udder cleanliness to account for the infectious nature of the disease has the potential to improve prediction accuracies in this context.

Prediction of ketosis works significantly better if the full data set is used (0.701 vs. 0.449). The high F1 score in the full data set is somewhat surprising, given there are only 673 observations of cows with ketosis in the data set. Existing studies on the prediction of metabolic status using ML approaches are mostly not directly comparable to our approach, as they either directly predict *β*-hydroxybutyrate levels^34, 35^, culling risk^36^ or cluster observations rather than predicting outcomes^37^. The only comparable study^38^ uses random forests and support vector machines to predict poor metabolic status. In this study, random forests achieved higher sensitivity (67.8-82.8%) but lower specificity (76.7-88.5%) than our approach (the support vector machines performed similarly). Along the same lines, prediction of periparturient hypocalcemia works considerably better if the full data set is used (F1=0.619 vs. F1=0.275). The stark difference between prediction performance using the full and the reduced data set for both metabolic diseases evidences the high influence of the time before calving on the metabolic fate of the cows. The moderately high prediction performance is somewhat surprising, given there were only 293 observations of periparturient hypocalcemia in the data set.

Prediction of ovarian cysts achieves moderate F1 scores of F1=0.482 and F1=0.464 in the full and the reduced data set respectively. Prediction of metritis works a bit better with F1 scores of F1=0.611 and 0.500 in the full and reduced data set, respectively. Again, the moderately successful prediction of metritis is surprising, given that there are only 265 observations of metritis in the data set. There are no other studies that try to predict the incidence of ovarian cysts or metritis in dairy cattle based on machine learning approaches that our results could be compared to.

For the remainder of this work, we will focus our analysis on the five diseases for which we achieved prediction performances with F1 scores ≥ 0.5: lameness, anoestrus, ketosis, periparturient hypocalcaemia and metritis. Since the random forests performed considerably better on the full data set, we use the full data set going forward. The exception to this is the analysis of anoestrus: a closer inspection of feature importances of the random forest using the full data set (see section 2.2 below for a more detailed description) revealed that the high prediction performance could almost exclusively be attributed to the features that reflect a farm-based bias in disease reporting (see sec. 4.1 for details). The classifier for the data set which only includes observations of lactating cows achieved a similarly high performance (F1=0.720 vs. F1=0.731 in the full data set) and only relies on reporting bias to a much smaller extent. Therefore, in the following analysis of anoestrus, we exclude observations from dry cows.

### 2.2 Feature importances and disease risk

In the following, we investigate the influence of feature groups and single features on the five diseases that achieved a prediction performance of at least F1=0.5. We calculate the permutation importance^52^ of all 138 features over 1,000 repeats. To give a first overview over feature importances, we grouped the features into the eleven categories that are illustrated in Fig. 1 (see sec. 4.1 for more details). We calculate the cumulative permutation importance of all features in a given category. Due to possible correlations between features, the sum of all permutation importances is not guaranteed to be one, so we normalise permutation importances by the sum of all permutation importances and report contributions to overall permutation importance in %. Results for the eleven feature groups and five diseases are shown in Tab. 3.

**Table 3.** Cumulative permutation feature importance contributions for the eleven feature categories and five diseases with a prediction F1 score of ≥ 0.5.

| feature category | lameness [%] | anoestrus [%] | ketosis [%] | periparturient hypocalcemia [%] | metritis [%] |
| --- | --- | --- | --- | --- | --- |
| housing | 28.2 | 30.8 | 23.4 | 12.5 | 10.6 |
| feed | 18.7 | 26.8 | 26.3 | 13.9 | 23.1 |
| environment | 11.1 | 6.7 | 12.2 | 2.5 | 11.4 |
| milk | 1.5 | 4.7 | 6.6 | 13.7 | 8.1 |
| lactation stage | 0.4 | 1.5 | 6.3 | 25.9 | 2.1 |
| husbandry | 12.7 | 13.1 | 8.6 | 4.2 | 4.7 |
| age | 10.9 | 8.7 | 1.0 | 19.8 | 2.4 |
| physique | 4.0 | 2.7 | 5.8 | 5.0 | 5.2 |
| breeding values | 1.9 | 4.1 | 3.3 | 1.9 | 3.3 |
| breed | 0.6 | 0.2 | 1.9 | 0.4 | 0.9 |
| diagnosis source | 9.9 | 8.7 | 4.6 | 0.3 | 28.5 |

In general, no single feature category dominates for any disease and two or more categories are usually necessary to explain more than 50% of the contributions to permutation importance to a disease. Housing and feed have the largest contributions to feature importance across all five diseases. Notably, environmental features – which up to this point have gotten relatively little attention in dairy cattle health research, evidenced by the lack of studies – have a moderately high contribution to all diagnoses except periparturient hypocalcemia. Milk- and lactation -related feature contributions show a large variance with contributions depending on disease type: on the high end, milk- and lactation-related features have a contribution of 13.7% and 25.9% respectively to periparturient hypocalcemia prediction feature importance and 6.6% and 6.3% to ketosis diagnoses. In addition, milk-related features also contribute 8.1% to permutation importance in metritis prediction. On the low end, milk- and lactation-related features have next to no contribution to lameness and anoestrus. Lameness and anoestrus are on the other hand considerably influenced by husbandry, as is ketosis. Age (which includes parity) plays a major role in the prediction of lameness (10.9%) and periparturient hypocalcemia (19.8%) and next to no role in the prediction of other diseases. Physical indicators such as BCS, play only a minor role for permutation importances for all diseases. Interestingly, breed also only has insignificant contributions to permutation importance, even though there is significant variance of breeds in the data set, with 3327 Fleckvieh, 1376 Braunvieh and 1061 Holstein animals. The diagnosis source (veterinarian, state control association employee, observation near culling or calving, or based on lameness score/ketosis test) has significant contributions to all diseases except for periparturient hypocalcemia. This is expected since farms seem to over- or under-report certain diagnoses based on their dominating diagnosis source.

On its own the permutation importance of a feature does not provide any information about the effect of a feature on the disease risk for a given disease. The permutation importance only provides information about the importance of the presence or absence of a feature (in case of a categorical feature) or high/low value of a feature (in case of a numerical feature) for the prediction of the disease. Nevertheless, a given feature can be associated with a decreased or increased risk for a disease. To investigate the *direction* of the association of a single feature to disease risk, we use multivariate logistic regression. Details of the logistic regression model are described in sec. 4.3. We report the permutation importances of the 50 most important individual features and their respective odds ratios for lameness, anoestrus, ketosis, periparturient hypocalcemia and metritis from the logistic regression in the supplement in tables 14, 15, 16, 17 and 18.

In the following, we discuss the importance for prediction and influence on disease risk of single features, focusing on features with high importance which are associated with significant reductions or increases of the odds of diseases. Our aim is to compare the effects we find for individual features with known effects in the literature and to highlight features that are of great importance in our analysis yet have not been addressed in the existing literature so far. We focus on the 50 most important features for each of the five diseases, including features that rank below 50 only in cases where they play a prominent role in the literature. All odds ratios reported in the following analysis have a significance level of at least *p* < 0.05, we report 95% confidence intervals in square brackets next to the odds ratios. The individual *p – values* for all features are listed in the corresponding tables in the appendix.

#### 2.2.1 Lameness

Lameness is characterised by an abnormal gait that is often caused by pain or discomfort. Lameness can be caused by a range of individual conditions, such as digital dermatitis, or other foot and claw disorders^53^. In the diagnosis used for this analysis, these individual conditions are summarised in an overarching “lameness” diagnosis, which is characterised by a lameness score, based on an assessment of animal mobility behaviour^54^ or a veterinarian’s diagnosis.

In a recent meta study^11^, Oehm et al. identify five robust risk factors for lameness: parity, body condition score (BCS), herd size, days in milk, and claw overgrowth. Of these risk factors we find parity (OR=2.25 [2.17; 2.33]), body condition score (OR=0.71 [0.68; 0.73]) and herd size (OR=1.16 [1.11; 1.21]) ranking highly among the most important features for disease prediction (see supplement table 14). Days in milk ranks relatively low with a feature importance of 0.12 ± 0.19% and claw overgrowth is not assessed in the available data. Odds ratios for BCS show a very similar association of high BCS with decreased disease risk (OR=0.71 [0.68; 0.73]) and an association of larger herd sizes with increased risk (OR=1.16 [1.11; 1.21]). This is also consistent with the association of good muscularity score – which is highly correlated with BCS (*R*^2^ = 0.59) – with decreased disease risk (OR=0.62 [0.60; 0.65]). On the other hand, large chest girth is associated with an increased risk for lameness (OR=1.26 [1.20;1.32]). Contrary to existing research^55, 56^, animal body weight decreases the odds for lameness (OR=0.84 [0.80; 0.89]), but does not rank highly among feature importances (rank 127). The strong association of parity on the risk for lameness in our analysis (OR=2.25 [2.17; 2.33]) lies within the confidence interval reported by the meta study (OR=1.63 [0.77; 3.46]).

In our analysis, a high yearly precipitation decreases the odds of lameness (OR=[0.80 [0.77; 0.84]) and mean relative humidity has a moderately detrimental effect (OR=1.09 [1.05; 1.14]). This is contrary to two reports about increased lameness of cows due to softer claws in high-moisture environmental conditions^57, 58^. We find that both a high standard deviation in temperature as well as a high mean yearly temperature increase the odds for lameness much stronger (OR=1.40 [1.34; 1.46] and OR=1.23 [1.18; 1.29], respectively). This is in line with research that suggests a connection between heat stress and increased standing time^13^, which is argued to be a major risk factor for lameness^59, 60^. Consistent with this finding, a high number of yearly low temperature days is associated with a decreased risk for lameness in our analysis (OR=0.89 [0.86; 0.93]). Somewhat counter-intuitively, a high number of high-wind days increases the odds of lameness (OR=1.18 [1.13; 1.22]).

Chopped straw litter in free-stalls for lactating and dry cows is associated with a significantly increased risk of lameness (OR=1.54 [1.41; 1.69] and OR=1.63 [1.49:1.79]) (consistent with a decrease in risk if other litter is used (OR=0.80 [0.72; 0.88])). As these features do not significantly correlate with other housing or husbandry features, that could explain the effect, the relation to lameness is not obvious to us. Similarly, slits in the stable floors and the absence of open-air areas increase the odds of for lameness across the board, in line with literature that finds increased prevalence of lameness in pigs kept on slatted floors^61^. Other important housing features are deep bed cubicles and deep litter cubicles, which are both associated with a significantly decreased disease risk for lameness (OR=0.39 [0.36; 0.43] and OR=0.56 [0.48; 0.64], respectively). Free-stall systems with deep-bed cublices have been reported to be associated with reduced risk for lameness before^62–65^.

Other housing features that are related to a reduced lameness risk are related to the milking system of the farm: the existence of an automated milking switch-off (OR=0.70 [0.63; 0.77]), pipe milking stalls (OR=0.47 [0.39; 0.56]) and a higher milking vacuum (OR=0.87 [0.84; 0.91]). Of these features, pipe milking stalls are moderately to strongly correlated with tie stalls for heifers (*R*^2^ = 0.62) and lactating cows (*R*^2^ = 0.76), which in turn reduce the odds for lameness (OR=0.64 [0.53; 0.78] and OR=0.71 [0.60; 0.83], respectively). This result is in line with literature^66, 67^ that reports a significantly reduced prevalence of lameness for cows kept in tie stalls vs. cows kept in free-stalls. A higher milking vacuum is weakly correlated with both pipe milking stalls (*R*^2^ = 0.22) and tie stalls for lactating cows (*R*^2^ = 0.21), which could explain why this feature is related to reduced odds for lameness. The existence of an automated milking switch-off is weakly correlated to “other” litter types in free-stalls for lactating cows (*R*^2^ = 0.20), dry cows (*R*^2^ = 0.20) and heifers (*R*^2^ = 0.18), which are related to reduced odds of lameness. In a previous study, hock lesions, an indicator for lameness, were associated with herringbone milking parlours^68^.

A high Total Merit Index is associated with a moderately decreased lameness risk (OR=0.78 [0.74; 0.81]), which is consistent with the fact that longevity, which is combined with the auxiliary trait feet and legs from conformation scoring, is a highly weighted trait in the TMI^69^. The fitness index itself, also including longevity, ranks low (145) but also reduced the odds for lameness (OR=0.71 [0.68; 0.74]. A high claw trimming frequency is associated with an increased lameness risk (OR=1.18 [1.06; 1.32]). Nevertheless, it is possible that in this case causality is reversed, since farms with lameness problems in their herds will prioritise claw trimming. Lastly, the annual herd milk yield average is an indicator for production intensity of a farm and is associated with a slightly decreased risk for lameness (OR=0.88 [0.84; 0.92]). Organic farms also have a comparatively lower incidence of lameness (OR=0.61 [0.54; 0.68]). Similarly, separated ration types of staple forage and concentrate feed, which are associated with a decreased risk for lameness (OR=0.65 [0.60, 0.71]) are possibly also an indicator for a less intense production, as corn silage is a known risk factor for lameness^70–72^.

#### 2.2.2 Anoestrus

Anoestrus is a diagnosis that indicates the absence of oestrus signs in an animal. This can be caused by both silent ovulation, i.e. the absence of oestrus signs alltogether, or by a failure in oestrus detection on the side of farm management^73^. In our data set, anoestrus diagnosis are a combination of all of these underlying causes that could lead to a failure of the cow of entering oestrus or a failure of the farmer to detect oestrus.

In accordance to the literature^74–76^, we find that a high production intensity, i.e. high milk yield, is associated with an increased risk for anoestrus. This can be explained by the inherently low expression of oestrus signs in high-production dairy cows^77^. A range of features in our analysis is associated with a high production intensity and shows associations with increased or decreased risk for anoestrus accordingly. Features associated with an increased risk are the test-day milk yield of the animal (OR=1.52 [1.37; 1.70]), annual herd milk yield average (OR=1.65 [1.50; 1.80]) and herd size (OR=1.37 [1.26; 1.48]). As oestrus is expected relatively early in the lactation of a cow, the association of a high number of days in milk with a low risk for anoestrus (OR=0.61 [0.56; 0.68]) is consistent. Similarly, the association of increased anoestrus risk with the season autumn (OR= 2.04 [1.76; 2.36]) is most likely associated to management choices and the pregnancy cycle of cows as cows kept on alpine pastures are expected to enter oestrus in autumn – but could also be related to increased heat stress during the summer months^78^. The contribution of breeding values to the prediction of oestrus is consistent with breeding targets and the relation of high milk yield to increased risk of anoestrus: the fitness index – which contains a fertility component – is associated with decreased risk of anoestrus (OR=0.76 [0.69; 0.83]) while the milk index is associated with an increased risk (OR=1.34 [1.23; 1.47]).

Estrus detection is frequently based on the observation of increased movement activity of cows^79, 80^. In this context, the strong association of the absence of open air areas for animals with an increased risk of anoestrus (OR=2.95 [2.48; 3.52] for lactating cows and OR=3.83 [3.08; 4.77] for dry cows) might be an indicator for increased difficulties in oestrus detection for cows that have limited opportunities for movement. Regarding animal housing, the strong association of outdoor climate housing with open fronts with increased disease risk (OR=3.20 [2.68; 3.82]) is noteworthy. This barn design is neither strongly correlated to high intensity farms (*R*^2^ = 0.21 with yearly milk yield per cow), nor to the absence of open-air areas for animals (*R*^2^=0.15 with the absence of open-air areas for lactating cows). There is no explanation for this association in the existing literature either.

There are a range of features relating to animal nutrition that play an important role in the prediction of anoestrus in cows. The emergent pattern shows that features related to an energy-rich nutrition are associated with an increased anoestrus risk, whereas features related to a high amount of crude fibres in rations are related to a decreased anoestrus risk. Specifically, the total amount of nitrogen free extracts (OR=1.96 [1.78; 2.17]), the content of metabolisable energy in rations (OR=1.66 [1.46; 1.88]), the content of utilisable protein (OR=1.45 [1.30; 1.62]), the total amount of undegraded dietary protein (OR=1.74 [1.58; 1.92]), the dietary proportion of concentrates (OR=1.42 [1.27; 1.60]) and the total amount of ether extracts (OR=1.44 [1.32; 1.157]) all are associated with an increased risk of anoestrus. Moreover, staple forage types that contain energy-rich ingredients in addition to grass silage, hay, grass and field forage silage are associated with an increased risk of anoestrus: corn silage (OR=1.72 [1.40; 2.13]) or corn (OR=2.29 [1.85: 3.07]). On the other hand, the total amount of crude fibre (OR=0.85 [0.77: 0.93]) and the total amount of crude ash (OR=0.65 [0.58; 0.72]) in rations is associated with a decreased risk of anoestrus, as is the time on pasture of lactating cows (OR=0.81 [0.62; 1.07]), the dietary proportion of grass silage (OR=0.61 [0.56; 0.67]) and partial mixed ration type for forage (OR=0.56 [0.46; 0.70]). In general, animals that mobilise a large amount of body mass at the beginning of lactation seem to be at higher risk for anoestrus^81, 82^. Under these conditions, the association of high-energy rations with increased risk for anoestrus might be interpreted as a reflection of an already occurring preventive management intervention to prevent anoestrus in at-risk cows. To clarify this relationship further, ration change protocols in farms in relation to the identification of animals at risk of metabolic or fertility disorders would need to be investigated in more detail. Findings in the literature about a relation between BCS and anoestrus are not always consistent^12^. Nevertheless, a range of studies reports an association of a decrease of risk of anoestrus with higher BCS^83, 84^, which is consistent with our findings (OR=0.83 [0.76: 0.90]).

There are very few reports about the influence of environmental factors such as precipitation, wind and ambient temperature on oestrus. One study reports that lower conception rates are associated with increased rainfall^78^, but it is questionable if conception rates can be directly associated with oestrus. In any case, we find that mean yearly precipitation is associated with a marked decrease in the risk for anoestrus (OR=0.50 [0.44; 0.57]). Temperature effects do not rank high in the feature importances and contradict the literature, which reports a positive correlation between oestrus activity^85^: yearly temperature is associated with a significantly increased risk of anoestrus (OR=2.99 [2.60; 3.43], p < 0.001, rank 111, not listed in table) and, accordingly, the number of low temperature days is associated with a decreased risk (OR=0.61 [0.54; 0.68], p < 0.001, rank 88, not listed in table). Nevertheless, the literature also reports a negative correlation of conception rate with ambient temperature^78^. There are no reports about the influence of wind on oestrus activity in the literature. In our analysis, the number of high wind days both ranks highly in the feature importances and is associated with an increased risk of anoestrus (OR=1.29 [1.19; 1.40]).

In our analysis, parity is associated with an increased risk for anoestrus (OR=1.45 [1.34; 1.58]). In the literature, there are conflicting reports about the relationship between parity and anoestrus as there are both reports of decreased oestrus intensity for multiparous cows^86^ as well as an increase in mounting behaviour with parity^85^.

#### 2.2.3 Ketosis

Ketosis is caused by a negative energy balance during high milk production, and is most likely to occur during the first weeks of lactation. A negative energy balance leads to an increased mobilisation of body fat, resulting in a ketone mobilisation that is higher than its utilisation. Symptoms of ketosis are reduced milk yield, loss of body weight, reduced appetite, fever and a high count of ketone bodies in the milk. Ketosis is divided in sublinical ketosis – a high count of ketone bodies without other symptoms – and clinical ketosis, where animals exhibit the previously described symptoms. In this analysis we combine both diagnoses into one ketosis label.

Some features related with a high-intensity production environment are associated with an increased risk of ketosis: large herd sizes increase the odds of ketosis OR=1.37 [1.26; 1.48], which is consistent with reports in the literature^87, 88^, while the annual herd milk yield average is not significantly associated with an increase or decrease of the odds of ketosis (OR=1.08 [0.98; 1.19]. Features related to the body condition of the cows rank highly among the important features and almost unanimously are positively associated with risk for ketosis (OR=2.38 [2.14; 2.64] for body weight, OR=1.96 [1.79; 2.14] for chest girth, OR=1.58 [1.43; 1.75] for waist circumference and OR=1.70 [1.56, 1.87] for BCS). This is consistent with ubiquitous reports about an association between high BCS and risk of ketosis^46, 50, 89–91^.

Parity ranks 51^th^ in the feature importance for prediction of ketosis and is associated with a moderately increased risk for ketosis (OR=1.45 [1.35; 1.57]). This is consistent with the literature which mostly reports increasing risk for ketosis with parity^50, 87, 92, 93^.

Two studies find an association between higher milk fat percentage and lower milk protein percentage^50, 91^ or high milk fat-protein ratio^46^ and an increased risk of clinical ketosis. Our analysis also finds an association between high milk fat percentage and ketosis risk (OR=1.54 [1.39; 1.70]) and high milk fat-protein ratio (OR=1.53 [1.40; 1.68], rank 170). In addition, our analysis finds that a high milk protein percentage is also related with an increased risk of ketosis (OR=1.43 [1.25; 1.65]). In the literature, high test-day milk yield is associated with an increased risk for ketosis^50, 94^, whereas our analysis finds a relation with a decreased risk (OR=0.80 [0.67; 0.97]). Reports on the relation of SCC and ketosis are scarce. One study finds no association of SCC with ketosis^50^, whereas we find an association of increased risk of ketosis with high SCC (OR=1.63 [1.46; 1.82]). We also report a moderate association of milk urea content with a decreased risk of ketosis (OR=0.89 [0.78; 1.02]).

Two studies report a negative association between high ketosis incidence and high ambient temperature-humidity index^95^ at calving or high temperature^87^ at calving, respectively. This contradicts our findings, which show an association of high mean yearly temperature (OR=1.23 [1.11; 1.35]), temperature deviation (Or=1.89 [1.71; 2.09]) and high mean relative humidity (OR=1.39; 1.26; 1.54) with increased clinical ketosis risk. To our knowledge, there are no reports of associations of other environmental parameters with ketosis in the literature. In our analysis, we find a marked association between precipitation and a decreased risk of ketosis (OR=0.46 [0.41; 0.52]) as well as the standard deviation of humidity (OR=0.80 [0.73; 0.87]).

Features related to cow energy intake are of great importance for ketosis prediction across the board. Most notably, all features related to a high energy and protein content in rations fed to the animals in the time before the diagnosis are associated with a decreased risk of ketosis (OR=0.57 [0.51; 0.63] for the dietary proportion of concentrates, OR=0.49 [0.44: 0.54] for the amount of crude ash in rations, OR=0.45 [0.41; 0.51] for the amount of crude fat in rations, OR=0.85 [0.78: 0.93] for the amount of nitrogen free extracts in rations, OR=0.48 [0.43; 0.54] for the amount of undegraded protein in rations, OR=0.64 [0.59; 0.70] for the amount of metabolisable energy in rations and OR=0.74 [0.68; 0.81] for the ruminant nitrogen balance). On the other hand, year-round feeding of corn silage is associated with an increased risk of ketosis (OR=1.43 [1.15; 1.77]), as is feeding of silages with a high energy content (OR=2.24 [1.86; 2.70]). In addition, a feed distribution system for high performance feed that is neither on demand nor exact allotment or manual allotment two times a day is associated with a marked increase in ketosis risk (OR=2.54 [2.07; 3.10]). The literature about the influence of concentrate feed intake in relation to ketosis is inconsistent, reporting both positive and negative associations of large amount of concentrate feed with ketosis risk^96^ and complicated relations between the management of feed intake and ketosis risk^97^. The information on rations used in this study only reports ration composition and not the amount of feed actually consumed by the animals. Therefore, to untangle the complicated inter-dependencies of nutrient intake and feed intake management, a more detailed recording of the actual feed consumed by the animals is probably needed.

Features related to animal housing are less significant for the prediction of ketosis than for other diseases and are also not reported in the literature. Nevertheless, a range of features are of importance and are associated with moderately decreased or increased risk of ketosis. Notably, two features related to the housing of young stock are associated with increased ketosis incidence: chopped straw as litter for calf free-stalls, lying mats in calf free-stalls, as well as a free-stall housing system that is not deep or high bed cubicles, deep litter or sloped floors (OR=2.56 [2.02; 3.25], OR=1.33 [1.08; 1.63], and OR=1.64 [1.32; 2.03], respectively). Chopped straw as litter in free-stalls for lactating cows is also associated with an increased risk (OR=1.80 [1.49; 2.17]). manure removal: slurry with solid flooring is associated with an increased risk for ketosis (OR=1.27 [1.08; 1.50]), whereas a walkway floor in the free-stall for dry cows that is neither solid concrete (with or without slits) nor rubber mats is associated with a decreased risk (OR=0.62 [0.49; 0.79]).

#### 2.2.4 Periparturient hypocalcemia

Postparturient hypocalcemia is a disease that is characterised by low blood calcium levels. Similar to ketosis, periparturient hypocalcemia occurs early in lactation when calcium demand for milk production is increased and results in restlessness, followed by an inability to stand and, finally, unconsciousness and death.

Consistent with ubiquitous reports in the literature^98, 99^, the by far most important feature for the prediction of periparturient hypocalcemia is parity, which is strongly related with an increase of periparturient hypocalcemia risk (OR=2.39 [2.23; 2.55]). This is due to the decreased ability of cows to absorb calcium with increasing age. In addition, we find that days in milk are also associated with an increased risk of periparturient hypocalcemia (OR=2.22 [2.06; 2.40]).

Periparturient hypocalcemia is caused by an imbalance between the availability and the demand for calcium. Accordingly, nutrition-related features play a key role in the prediction of periparturient hypocalcemia: in line with recent observations in the literature^100^, the total amount of crude fibre in rations (OR=0.70 [0.64; 0.75]) is associated with a decreased risk of periparturient hypocalcemia. This is also true for the total amount of undegraded protein (OR=0.63 [0.56; 0.70]) and crude protein (OR=76 [0.70; 0.82]) are associated with a decreased risk of periparturient hypocalcemia. Furthermore, on-demand high performance feed distribution and partial mixed rations are also associated with decreased odds (OR=0.70 [0.59; 0.84] and OR=0.79 [0.65; 0.96], respectively), while total mixed rations are associated with increased odds (OR=1.83 [1.33; 2.51]), consistent with findings in literature that report detrimental effects of total mixed rations on animal welfare^101^. On the other hand, features related to a high milk energy content are associated with increased periparturient hypocalcemia risk across the board (milk protein percentage OR=1.26 [1.09; 1.45], milk fat percentage OR=1.15 [1.01; 1.30] and test-day milk yield OR=1.24 [1.04; 1.49]). The association of high milk yield with increased risk of periparturient hypocalcemia is consistent with reports in literature^43^. We also find an association of decreased risk of periparturient hypocalcemia in organic farms (OR=0.70 [0.55; 0.89]), which can be explained by the high negative correlation of milk yield with organic farm type (*R*^2^ = −0.41).

Features associated with an obese body condition are also associated with increased risk of periparturient hypocalcemia. Odds ratios range from 1.24 [1.14; 1.36] for BCS over OR=1.27 [1.16; 1.40] for chest girth to OR=1.70 [1.53; 1.88] for animal mass and OR=1.79 [1.62; 1.97] for waist circumference. A relation between high BCS and increased risk of periparturient hypocalcemia is also reported in the literature^12, 43, 102^.

Many features related to housing rank high in the feature importance for the prediction of periparturient hypocalcemia. As with other diseases except for lameness, literature on the influence of different housing parameters on periparturient hypocalcemia is next to non-existant. In our analysis, we find that chopped straw as litter in free-stalls for lactating cows (OR=1.26 [1.05; 1.51]), dry cows (OR=1.42 [1.18; 1.72]) and young stock (OR=1.94 [1.52; 2.48]) is consistently associated with an increased risk of periparturient hypocalcemia. On the other hand, long-straw litter in free-stalls for dry cows is associated with a slightly reduced risk for periparturient hypocalcemia (OR=0.87 [0.70; 1.08]). High-bed cubicles in free-stalls for dry cows are associated with an increased risk (OR=[1.44 [1.11; 1-88]]), as is manure removal through slits (OR=1-54 [1.27; 1.87]) and a high number of milking places (OR=1.13 [1.0; 1.23]). On the other hand, milking robots are associated with a decreased risk (OR=0.63 [0.49; 0.83]).

Environmental conditions play a secondary role in the prediction of periparturient hypocalcemia. The association of a high standard deviation of temperature with increased risk of periparturient hypocalcemia (OR=1.48 [1.34; 1.63] is consistent with a similar finding in the literature^99^. On the other hand, contrary to that study we find a relation of decreased risk of periparturient hypocalcemia with mean precipitation (OR=0.77 [0.69; 0.85]). Another recent study among a large number of cows that considered the ecological life zone^103^ unfortunately does not make any statement about the direct relationship of temperature, humidity or precipitation with periparturient hypocalcemia.

#### 2.2.5 Metritis

Metritis is an inflammation of the uterus walls shortly (within 10-21 days) after calving. Symptoms include an enlarged uterus and purulent discharge, leading to reduced milk yield and fever and, ultimately, collapse and death if untreated.

The literature reports either an u-shaped relation between parity and metritis^104^, with an association of increased risk for metritis in heifers and third-parity and above cows or a decreasing risk of metritis with parity^48^. Our linear-regression based approach of qualifying the influence of features on disease risk does not resolve non-linear dependencies and we find an association of parity with increased risk (OR=1.64 [1.49; 1.81]), somewhat contradicting results in literature. Literature also reports increased odds for metritis in cows calving between November and April. Our analysis is somewhat consistent with these findings, with increased odds for metritis in autumn (OR=2.38 [1.90; 2.98]), decreased odds for births in spring and summer (OR=0.71 [0.53; 0.94] and OR=0.51 [0.37; 0.70], respectively) and inconclusive results for winter.

Consistent with the literature^105^ we find no significant influence of BCS on the odds of metritis. On the other hand, we find that high body weight (OR=1.42 [1.22; 1.66]), waist circumference OR=1.17 [1.01; 1.36]) and chest girth (OR=1.38 [1.21; 1.58]) all increase the odds of metritis. One existing study shows a relation between increased dry matter intake, feeding time and reduced odds for metritis. This could be related to the strongly decreased odds of metritis for separate concentrate feed ration types (OR=0.35 [0.26; 0.46]), as this feeding management strategy possibly reduces aggressive interactions between cows at feed banks and therefore increases feeding time. In addition, we find a range of feed-related features that decrease the odds of metritis: dietary proportion of concentrates (OR=0.84 [0.73; 0.98]), total amount of crude fibre in the ration (OR=0.74 [0.65; 0.84]), total amount of crude protein in the ration (OR=0.84 [0.74; 0.96]), total amount of usable protein in the ration (OR=0.83 [0.72; 0.96]), total amount of crude ash in the ration (OR=0.75 [0.66; 0.86]) and total amount of undegraded protein in the ration (OR=0.85 [0.74; 0.98]). Only the dietary proportion of straw is related to an increase in the odds for metritis (OR=1.40 [1.10; 1.79]). Regarding milk-related features, low protein percentage and milk urea content are associated with a slight reduction in the odds for metritis (OR=0.88 [0.74; 1.05] and OR=0.85 [0.73; 0.99], respectively).

As with other diseases, most of the existing studies do not include herds kept under different housing conditions and, therefore, cannot assess housing conditions as risk factor. In our analysis, we find a range of housing conditions for lactating and dry cows that are associated with metritis: chopped straw as litter in free-stalls increases the odds of metritis (OR=2.42 [1.81; 3.23], as does the absence of an open-air area for dry cows (OR=3.12 [2.33; 4.18]). Manure removal in dry cow free-stalls by scraper decreases the odds of metritis (OR=49 [0.38; 0.65]) whereas manure removal which was neither through slatted floors nor by scraper increased the odds (OR=1.41 [1.10; 1.80]). These findings could be related to the assumed cleanliness of the barn, yet one of the few studies that investigated housing effects on metritis did not find any effect of bedding on metritis risk^106^. Possibly along the same lines, a claw trimming frequency of twice per year is related to decreased odds for metritis (OR=0.57 [0.45; 0.73]).

To our knowledge, there are no existing studies that relate environmental conditions such as temperature or precipitation to risk of metritis. In our analysis we find that a high standard deviation of temperature increases the odds for metritis (OR=2.94 [2.45; 3.53]), whereas a high number of low temperature days as well as high yearly precipitation decrease the odds (OR=0.67 [0.57; 0.78] and OR=0.59 [0.49: 0.73], respectively). Lastly, we also find a moderate association between the Total Merit Index and a decrease in the odds for metritis (OR=0.85 [0.75; 0.98]).

## 3 Conclusion

Here, we have demonstrated the value of combining different data sources from various life domains of dairy cattle for the prediction of common metabolic, locomotive and reproductive diseases of cows. We apply a random-forest based approach to predict occurrences of lameness, acute and chronic mastitis, anoestrus, ovarian cysts, metritis, periparturient hypocalcemia and ketosis. Our data set consists of 22,923 observations from 5,828 animals from 166 different herds, including 138 different features from 11 different animal life domains. Life domains include animal breed, age, lactation stage and physique as well as nutrition, milk parameters, breeding values, housing and husbandry and environmental factors and, as target variable, diagnoses. To interpret the importance of individual features to disease prediction, we use feature permutation importances in combination with multivariate logistic regression. We achieve considerable success in predicting lameness, anoestrus, periparturient hypocalcemia, ketosis and metritis, with F1 scores between F1=0.611 and F1=0.72. Interpretation of feature contributions uncovers the complex interactions and multiple contributions of different life domains. We confirm the association of a large number of features with increased odds of diseases reported in the literature and report associations between a range of features and disease risk that have never been reported before. With this work we hope to show that the integration of data from multiple sources can create added value in precision livestock farming and that outcomes can be used to improve animal wellbeing.

## 4 Methods

### 4.1 Data

The data used for the prediction of disease risks consisted of 22,923 observations from dairy cows collected in the time between January 2 2014 and January 30 2016. Here, the additional features related to feed intake and animal physique were only observed during the year 2014 within the scope of the Efficient Cow project^46^, while other features were available for the whole period. These observations stem from 5,828 dairy animals that were kept at 166 farms. Observations have specified dates and the same animal can contribute observations from times at which it was healthy as well as observations associated with a diagnosis. Every observation has 138 features (87 numerical, 51 categorical) from 11 feature categories (illustrated in fig. 1, which were aggregated from 6 different sources: the national breeding registry, the national cattle disease registry, a farm survey, recurring dairy herd improvement (DHI) assessments extended by assessments and records from the national weather service. A list of all features is given in the supplement, tables 3 to 13.

**Diagnoses** and **diagnosis origin** (supplement table 3): An observation contains one out of 56 possible diagnoses or refers to a cow that was healthy at the time of observation. A total of 9,260 observations have a diagnosis (40.4%) while 13,663 observations belong to healthy animals (59.6%). The eight most frequent diagnoses comprise 88.5% of all diagnoses and their frequency is listed in table 1. Diagnoses were taken from a national registry of cattle diagnoses^48^ together with a diagnosis date. We enriched these diagnoses by additional diagnoses for lameness from lameness scoring^53, 54^. Cows with a lameness score ≥ 3 (“moderately lame”) were given a “lame” diagnosis. We chose this threshold value to achieve a good contrast between only mildly lame cows (that are considered healthy in this approach) and severely lame cows. For the future, including information about lameness scores in the prediction as the potential to make the prediction more accurate. Next to lameness, we also enriched the diagnoses with additional ketosis diagnoses based on ketosis tests. To detect ketosis with a high sensitivity and specificity^107^, cow blood was tested using Keto-Tests 7 days and 14 days after calving. Cows with an amount of beta-hydroxybutyrate, a ketone, of ≥ 200*μ*mol/l in one or both tests were given a “ketosis” diagnosis. Diagnoses of periparturient hypocalcemia include both observations close to calving as well as diagnoses by veterinarians. Therefore, these diagnoses are a combination of diagnoses based on the observation of symptoms and diagnoses based on measurements of calcium levels in blood tests.

Entries in the national diagnosis registry can have different sources and different ways to be entered in the registry: entries can either be diagnoses by a veterinarian or observations at calving or a culling reason, observed by a farmer. Diagnoses can then be entered into the registry either automatically by veterinarians through an easy-to-use interface or by hand by an employee of the national performance recording organisation. These different origins and ways to enter information into the registry introduced a large bias into our data set, as the ratio of different diagnosis sources varies widely between farms and farms seem to over- or under-report different diagnoses, depending on what reporting pathway they rely most on. We therefore decided to introduce new features for the ratio of different diagnosis sources and their registry entering pathways at a given farm to capture the variance introduced by biased diagnoses sources.

**Housing** (supplement table 4): These 25 farm-specific features were collected once through a survey sent to all farms within the scope of the Efficient Cow project in Austria^46, 108^ and do not change over the course of time. Features include information for example on the number of cows, the means of manure removal and the type of flooring used in the stables. In a second study^109^, these features were used to create disease risk profiles of farms based on the living conditions. Some of the values within the features were very rare. These values were merged into an “other” value if there were fewer than 10 farms with a given value. **Husbandry** (supplement table 5): These 25 farm-specific features were collected through the same survey as described above. Features include information for example about whether or not a farm is organic, if and how long cows are grazing or on an alpine pasture and what type of milking system the farm has. Similar to the housing category, rare feature values were merged into an “other” value.

**Physical indicators** (supplement table 6): during recurring DHI assessments expanded by body and conformation assessments^110^, muscularity score, waist circumference, BCS^111^, animal body mass and chest girth of the animals were recorded as well.

**Milk indicators** (supplement table 7): At each test-day during the routine DHI assessments^110^, milk yield as well as fat, protein, lactose and urea content content and somatic cell count were acquired. Based on these records, fat and protein percentage, fat-protein ratio and energy corrected milk are routinely calculated and provided.

**Feed** (supplement table 8): These 40 features were also acquired as an extension of the of the recurring DHI assessments within the scope of the Efficient Cow project^46, 47, 110, 112^. They contain detailed information about the type of staple forage, ration type and the nutrient content of the rations (protein content, crude fibre content, …). Several measures such as the total organic mass of a ration were excluded since they were highly correlated (*R*^2^ > 0.9) with other measures and therefore considered redundant. Removing these redundant measures also slightly improved prediction accuracies (on the order of 0.01 F1). In addition, information about the quality of the feed was collected by assessors during farm visits: assessors rated the contamination of feed with mould its temperature (an indication for fermentation processes in the feed) on a scale from 1 (perfect) to 4 (insufficient). We used the average of these two ratings to create a binary indicator for feed quality: if the average rating was ≥ 1.1 (true for approximately 20% of farms), the indicator variable was set to “problematic feed quality”.

**Age** (supplement table 9): This category only includes two features: the animal’s parity and age at first calving.

**Lactation stage** (supplement table 10): Features related to the lactation category are days in milk, days pregnant and whether or not the animal was lactating at observation time.

**Breed** (supplement table 11): The data set also includes information about the main breed of an animal and ratio and type of any foreign genes^112^. This information stems from the national breeding registry^48^.

**Breeding values** (supplement table 12): Next to the genetic information we also include five breeding values from the same national breeding registry^113^: the Total Merit Index, milk index, beef index, fitness index and milk yield breeding value.

**Environment** (supplement table 13): Records from the national weather service from the year 2014 were used to calculate weather indicators for each farm based on its location. Weather indicators included in the data set are: the fraction of days with high wind conditions, i.e. at least one measurement on a given day with wind speeds > 2.5 m/s. The fraction of days with low temperature conditions, i.e. at least one measurement on a given day with a temperature < 0.5°C (cutoff value extracted from West 2003^114^). The yearly average temperature and temperature standard deviation, the yearly average precipitation and precipitation standard deviation, the yearly average relative humidity and relative humidity standard deviation, the altitude of the farm’s location and the season at the time of the observation. Season information is constructed from the observation date of the DHI assessment.

Data was aggregated using unique animal and farm IDs. Since observations of the DHI assessment and the body and conformation assessment did not necessarily occur at the same date as the diagnoses, diagnoses observations were matched to the nearest *preceding* DHI and body and conformation assessment observations and only diagnoses where the time difference between diagnosis and assessment was not larger than 365 days were kept (resulting in the number of diagnoses listed in tab. 1. Therefore, it does not make sense to use observations that occurred after a diagnosis. In addition to the data set matching, we also removed a range of features with a high correlation (*R*^2^ > 0.9) to another feature, or features which were just derivations of existing features such as the somatic cell count and its logarithm. For both the random-forest-based analysis and the logistic regression categorical features were one-hot encoded.

### 4.2 Prediction algorithm

Balanced random forest classifiers^115, 116^ were trained on the data for each of the eight most prevalent diagnoses and for two different versions of the data set: once on the full data set and once on the data set excluding observations from cows in the dry period. Before training, a holdout sample of ≈ 10% of observations was saved for final validation of the classifiers and removed from the training data set. The holdout observations were selected by first randomly selecting 6% of unique animals and then assigning all observations belonging to these animals to the holdout data set, amounting to approximately 10% of observations. This was done to prevent information leakage between the holdout data and the training data. We tried the same procedure but instead of randomly selecting animals, we randomly selected 20% of the farms and then assigned all cows and their observations belonging to these farms to the holdout data set. This drastically reduced the prediction quality, since there are a lot of different possible combinations of farm features, and the classifiers under-performed on farms for which they had not seen farms with similar management and living conditions during the training. Nevertheless, this approach could become feasible in the future, if a higher number of farms is included in the data set and every farm type is present in the training data set. Before training the random forest classifiers, missing values for features were imputed by the feature’s mean value.

To find the best hyper-parameters for the random forest classifiers, as a first step a randomised grid search over a large range of parameter values was performed. Parameters that were scanned were (i) the number of estimators in the forest (ii) the minimum number of samples for a node split (iii) the minimum number of samples in a leaf node (iv) the maximum depth of the trees (v) the maximum number of features in a tree and (vi) whether or not bootstrapping was applied. The randomised grid search was followed by a refined grid search for a smaller range of parameters around the resulting best parameter set from the randomised grid search. Both the randomised grid search and the refined grid search were performed with 10-fold cross-validation. The optimal parameters for each of the eight are listed in the supplement, table 2. The prediction accuracy of all classifiers was then tested on the holdout data set. Training and test accuracies and F1 scores are reported in Tab. 2.

### 4.3 Logistic regression

We apply multivariate logistic regression to investigate the individual effects of single features on diseases. These results provide the possibility to investigate the influence of a feature on the increase or reduction of the odds of a disease, rather than the pure *importance* of a feature for prediction. A logistic regression is a common method to investigate diseases risks. The dependent variable is the disease, which is explained by independent variables. The control group for every disease are all observations that were not matched to any other diagnosis. To minimise the effects of confounding variables in the regression model we adjusted for a number of important variables: parity, season, breed, diagnosis source well as the performance level of the farm. A disease risk was described as a function of the adjusted variables and the individual independent variables. We computed the individual risk for each feature–disease pair. Features with less then 10 observations or only observations in less then two herds in both the reference and the disease group were excluded.

## Acknowledgements

This work was conducted within the COMET-Project D4Dairy (Digitalisation, Data integration, Detection and Decision support in Dairying, Project number: 872039) that is supported by BMK, BMDW and the provinces of Lower Austria and Vienna in the framework of COMET-Compentence Centers for Excellent Technologies. The COMET program is handled by the FFG, grant number 872039.

The study was supported by the project “Efficient Cow” funded by the Austrian Federal Ministry of Agriculture, Regions, and Tourism (Vienna); the Federations of Austrian Fleckvieh (Zwettl), Brown-Swiss (Innsbruck), Holstein (Leoben) and the Federation of Austrian Cattle Breeders (Vienna), grant number 100861 BMLFUW-LE.1.3.2/0083-II/1/2012.

We thank the Zentralanstalt für Meteorologie and Geodynamik (ZAMG) Austria for supplying climate information for farm locations.

## Author contributions statement

J.L. Conducted the random forest analysis and drafted the manuscript.

C.M. Conducted the logistic regression analysis and revised the manuscript.

C.E.D. Initiated the D4Dairy project in general and contributed to the conceptualisation of this specific project within D4Dairy, organised funding of this project, provided data access, provided literature reviews and domain knowledge on dairy cow husbandry and contributed to revising the manuscript.

B.F.W. Contributed to the general idea of the D4Dairy project, provided literature reviews and domain knowledge on dairy cow husbandry, provided funding and data access to climate data and contributed to revising the manuscript.

F.S. Selected and provided data and contributed to revising the manuscript.

T.W. Contributed to the general idea of the D4Dairy project, provided literature reviews on metabolic and reproductive diseases, provided funding and contributed to revising the manuscript. P.K. Conceptualised the project, provided funding and data access and revised the manuscript.

## Additional information

The authors declare no competing interests.

