## Supplementary information for "Integrating diverse data sources to predict disease risk in dairy cattle"

| category | # features | source | examples | detailed description |
|---|---|---|---|---|
| diagnoses and diagnosis origin | 5 | national diagnoses registry[1], lameness scores, ketosis tests | lameness, ketosis, ovarian cysts | Tab. 3 |
| housing | 23 | farm survey[2] | stable type, manure removal system | Tab. 4 |
| husbandry | 23 | farm survey[2] | frequency of claw cleaning, pasturing of cows | Tab. 5 |
| physical indicators | 5 | extended DHI[3] | body condition score, weight | Tab. 6 |
| milk indicators | 8 | DHI[3] | somatic cell count, lactose content | Tab. 7 |
| feed | 40 | extended DHI[4] | ratio of concentrated feed, fodder contamination | Tab. 8 |
| breed | 14 | national cattle database (RDV)[1,5] | main breed, ratio and type of foreign genes | Tab. 11 |
| lactation stage | 3 | DHI[3] | days in milk, days pregnant | Tab. 10 |
| age | 2 | DHI[3] | parity, age at first calving | Tab. 9 |
| breeding values | 5 | national cow registry[6] | fitness score, TMI | Tab. 12 |
| environment | 10 | national weather service (ZAMG) | mean temperature, altitude | Tab. 13 |

**Table 1.** Feature categories and data sources of the data used in the training of the random forest classifiers.

| data set | diagnosis | # estimators | min split | min leaf | max depth | max features | bootstrap |
|---|---|---|---|---|---|---|---|
| full | lameness | 100 | 2 | 3 | 10 | sqrt | True |
| full | acute mastitis | 400 | 5 | 5 | 7 | log2 | False |
| full | anoestrus | 280 | 6 | 2 | 5 | log2 | True |
| full | ovarian cysts | 300 | 5 | 3 | 5 | log2 | False |
| full | periparturient hypocalcemia | 220 | 2 | 2 | 28 | log2 | False |
| full | ketosis | 280 | 2 | 1 | 35 | log2 | True |
| full | chronic mastitis | 400 | 5 | 4 | 5 | log2 | False |
| full | metritis | 420 | 6 | 3 | 5 | log2 | False |
| w/o dry | lameness | 300 | 3 | 3 | 11 | sqrt | True |
| w/o dry | acute mastitis | 400 | 6 | 4 | 5 | log2 | False |
| w/o dry | anoestrus | 300 | 4 | 1 | 7 | log2 | False |
| w/o dry | ovarian cysts | 300 | 4 | 3 | 5 | log2 | False |
| w/o dry | periparturient hypocalcemia | 380 | 6 | 3 | 5 | log2 | False |
| w/o dry | ketosis | 320 | 6 | 1 | 5 | log2 | False |
| w/o dry | chronic mastitis | 300 | 6 | 3 | 5 | log2 | False |
| w/o dry | metritis | 420 | 5 | 3 | 5 | log2 | False |

**Table 2.** Optimal parameters for the random forest classifiers of the eight most prevalent diseases, using two different versions of the data set.

**diagnoses**

| **feature** | **type** | **values** | **% missing** |
|---|---|---|---|
| diagnosis | categorical | healthy 59.60%, milk fever 2.94%, ketosis 2.87%, uterus inflammation 1.39%, anestrus 3.50%, acute mastitis 4.52%, chronic mastitis 1.46%, lameness 16.01%, other 7.71% | 0.00% |
| diagnosis source: culling reason or observation at calving | numerical | 0.24±0.21 % | 0.00% |
| diagnosis source: veterinarian | numerical | 0.22±0.26 % | 0.00% |
| diagnosis source: performance recording organisation (LKV) | numerical | 0.09±0.21 % | 0.00% |
| diagnosis source: score | numerical | 0.46±0.26 % | 0.00% |

**Table 3.** Features summarised in the feature category *diagnoses*: For categorical and binary features, the frequency of each category is reported. For numerical features, mean values and standard deviations are reported. For all features, the percentage of missing values in the data set is reported.

| housing | | | |
|---|---|---|---|
| **feature** | **type** | **values** | **% missing** |
| cubicle housing system | categorical | other 7.68%, deep bed cubicles and solid floors 48.31%, high bed cubicles and solid floors 9.78%, high bed cubicles and slatted floors 8.61%, deep bed cubicles and slatted floors 25.62% | 0.00% |
| manure removal | categorical | mixed forms 16.11%, slurry with perforated flooring 34.43%, solid manure 8.57%, slurry with solid flooring 40.89% | 0.00% |
| flooring in open air area for young stock | categorical | no open-air areas 55.51%, unpaved 4.00%, solid concrete 30.07% | 10.00% |
| floor in walkway of free-stall for young stock | categorical | other 19.62%, solid concrete 25.61%, concrete slits 36.76%, solid concrete with slits 4.30% | 14.00% |
| litter in free-stall for young stock | categorical | other 14.76%, chopped straw 20.66%, long straw 20.31% | 44.00% |
| manure removal in free-stall for young stock | categorical | slits 34.86%, scraper 20.53%, other 34.28% | 10.00% |
| free-stall system for young stock | categorical | other 14.32%, deep bed cubicle 17.29%, sloped floor 5.56%, high bed cubicle 33.44%, deep litter 18.06% | 11.00% |
| floor in open-air area for lactating cows | categorical | solid concrete 44.94%, other 10.88%, no open-air areas 43.17% | 1.00% |
| walkway floor in free-stall for lactating cows | categorical | solid concrete with slits 9.37%, solid concrete 20.65%, rubber mats 13.91%, rubberised slits 9.69%, concrete slits 16.97%, other 21.30% | 8.00% |
| litter in free-stall for lactating cows | categorical | chopped straw 46.97%, other 26.22%, long straw 15.96% | 11.00% |
| manure removal in free-stall for lactating cows | categorical | other 23.89%, scraper 40.49%, slits 28.86% | 7.00% |
| free-stall system for lactating cows | categorical | deep bed cubicle 72.04%, other 8.34%, high bed cubicle 13.02% | 7.00% |
| floor in open-air areas for dry cows | categorical | other 7.94%, solid concrete 34.86%, no open-air areas 53.03% | 4.00% |
| walkway floor in free-stall for dry cows | categorical | solid concrete with slits 11.63%, rubber mats 7.72%, other 22.76%, solid concrete 24.66%, concrete slits 19.75% | 13.00% |
| litter in free-stall for dry cows | categorical | long straw 17.98%, other 15.97%, chopped straw 43.64% | 22.00% |
| manure removal in free-stall for dry cows | categorical | slits 24.33%, other 32.24%, scraper 33.28% | 10.00% |
| free-stall system for dry cows | categorical | high bed cubicle 12.61%, other 15.05%, deep bed cubicle 51.30%, deep litter 11.27% | 10.00% |
| type of milking stalls | categorical | side-by-side 11.68%, tandem type 21.91%, herringbone parlour 41.98%, milking robot 12.53%, pipe milking 8.40% | 3.00% |
| silo type | categorical | silage bales 18.19%, no silo 8.21%, bunker silo 69.26% | 4.00% |
| barn design | categorical | outdoor climate house open front 18.78%, free-stall barn 44.03%, other 8.97%, outdoor climate house closed 22.96%, tie stall facility with pasture 5.27% | 0.00% |
| lying mats in free-stall for young stock | categorical | True 39.3%, False 32.2% | 28.5% |
| lying mats in free-stall for lactating cows | categorical | True 18.1%, False 55.4% | 26.4% |
| lying mats in free-stall for dry cows | categorical | True 19.7%, False 42.5% | 37.9% |

**Table 4.** Features summarised in the feature category *housing*: For categorical and binary features, the frequency of each category is reported. For numerical features, mean values and standard deviations are reported. For all features, the percentage of missing values in the data set is reported.

**husbandry**

| feature | type | values | % missing |
|---|---|---|---|
| claw trimming frequency | categorical | twice per year 56.70%, once per year 18.76%, three times per year 15.55%, only for lame animals 5.47% | 4.00% |
| milking unit removal | categorical | none 52.32%, present, including post-milking technology 10.94%, present 36.74% | 0.00% |
| dry cows group management | categorical | separate 53.62%, with lactating cows 38.58%, with young stock 7.74% | 0.00% |
| young stock on alpine pasture | categorical | True 41.4%, False 58.6% | 0.0% |
| lactating cows on alpine pasture | categorical | True 11.6%, False 88.4% | 0.0% |
| dry cows on alpine pasture | categorical | True 12.9%, False 87.1% | 0.0% |
| farm organically managed | categorical | True 19.9%, False 80.1% | 0.0% |
| young stock are kept in a tie stall facility | categorical | True 5.9%, False 91.4% | 2.7% |
| claw trimming done by farmer | categorical | True 73.7%, False 25.9% | 0.4% |
| lactating cows are kept in a tie stall facility | categorical | True 7.5%, False 92.5% | 0.0% |
| automated milking switch-off | categorical | True 80.5%, False 18.7% | 0.7% |
| milking stimulation | categorical | True 69.3%, False 30.7% | 0.0% |
| dry cows kept in tie stall facility | categorical | True 9.7%, False 90.2% | 0.1% |
| pasture of young stock | categorical | True 65.7%, False 34.3% | 0.0% |
| pasture of lactating cows | categorical | True 35.5%, False 64.5% | 0.0% |
| pasture of dry cows | categorical | True 36.1%, False 63.9% | 0.0% |
| young stock, time on pasture | numerical | 20.7±6.3 days | 41.83% |
| lactating cows, time on pasture | numerical | 7.5±5.23 days | 66.20% |
| pasture duration of dry cows | numerical | 15.0±9.1 days | 66.62% |
| number of milking places | numerical | 7.13±7.34 n | 3.93% |
| milking vacuum | numerical | 42.1±5.0 kPa | 7.80% |
| annual herd milk yield average | numerical | 8728.0±1528.0 kg | 0.00% |
| herd size | numerical | 40.2±18.8 cows | 0.00% |

**Table 5.** Features summarised in the feature category *husbandry*: For categorical and binary features, the frequency of each category is reported. For numerical features, mean values and standard deviations are reported. For all features, the percentage of missing values in the data set is reported.

**physique**

| feature | type | values | % missing |
|---|---|---|---|
| waist circumference | numerical | 256.0±16.0 cm | 0.71% |
| body condition score | numerical | 3.24±0.61 | 0.64% |
| muscularity score | numerical | 5.21±1.52 | 1.41% |
| chest girth | numerical | 210.0±12.0 cm | 0.69% |
| body weight | numerical | 701.0±97.0 kg | 0.35% |

**Table 6.** Features summarised in the feature category *physique*: For categorical and binary features, the frequency of each category is reported. For numerical features, mean values and standard deviations are reported. For all features, the percentage of missing values in the data set is reported.

| milk | | | |
|---|---|---|---|
| **feature** | **type** | **values** | **% missing** |
| test-day energy corrected milk daily yield | numerical | 34.2±23.5 kg | 20.21% |
| test-day protein yield percentage | numerical | 3.51±0.4 % | 18.88% |
| test-day fat-protein ratio | numerical | 1.29±0.25 | 93.42% |
| test-day fat yield percentage | numerical | 4.23±0.75 % | 18.88% |
| test-day urea content | numerical | 21.2±8.5 mg/dl | 19.05% |
| test-day lactose content | numerical | 4.75±0.21 % | 27.95% |
| test-day milk yield | numerical | 26.8±9.2 kg | 19.88% |
| test-day somatic cell count | numerical | 170329.0±466553.0 cells/ml | 18.94% |

**Table 7.** Features summarised in the feature category *milk*: For categorical and binary features, the frequency of each category is reported. For numerical features, mean values and standard deviations are reported. For all features, the percentage of missing values in the data set is reported.

**feed**

| feature | type | values | % missing |
|---|---|---|---|
| used forage types in diet | categorical | field forage silage, grass silage, hay, corn silage 16.16%, field forage silage, grass silage, hay, corn silage, pasture 14.11%, grass silage, hay 6.63%, grass silage, hay, pasture 7.67%, other 42.04% | 13.00% |
| forage | categorical | sequentially fed forages 54.54%, partial mixed ration 32.74% | 13.00% |
| forage type | categorical | year-round silage (with corn silage) 14.09%, other 11.46%, grass and grass products plus corn only 7.07%, grass plus corn and grass products plus corn 8.04%, grass and grass products only 19.51%, mixed ration with concentrates 39.84% | 0.00% |
| feeding group | categorical | lactating and dry cows on same forage (mixture) 35.01%, lactating cows 53.91%, dry cows 8.65%, cows on alpine pasture 1.19% | 1.00% |
| provision of concentrates | categorical | manual 12.94%, exact 77.60% | 9.00% |
| provision of supplementary concentrate | categorical | electronic feeder 70.44%, other 13.29%, exact 7.16%, manual, two times a day 9.11% | 0.00% |
| ration type | categorical | partial mixed ration 41.67%, feedstuffs sequentially fed 49.21%, total mixed ration 5.99% | 3.00% |
| problematic feed quality | categorical | True 15.5%, False 56.3% | 28.2% |
| dietary proportion of grass silage | numerical | 0.51±0.31 % | 0.38% |
| dietary proportion of green forage | numerical | 0.08±0.23 % | 0.38% |
| dietary proportion of hay | numerical | 0.14±0.22 % | 0.38% |
| dietary proportion of clover | numerical | 0.05±0.16 % | 0.38% |
| dietary proportion of concentrates | numerical | 0.25±0.14 % | 0.38% |
| dietary proportion of lucerne | numerical | 0.01±0.07 % | 0.38% |
| dietary proportion of corn silage | numerical | 0.19±0.21 % | 0.38% |
| dietary proportion of straw | numerical | 0.02±0.04 % | 0.38% |
| dietary proportion of cereal | numerical | 0.0±0.04 % | 0.38% |
| diet: total amount of acid detergent fibers | numerical | 0.0±0.0 g | 0.00% |
| diet: total amount of acid detergent lignin | numerical | 0.0±0.0 g | 0.00% |
| diet: total amount of neutral detergent fibers | numerical | 0.0±0.0 g | 0.00% |
| diet: total amount of net energy | numerical | 118.0±27.0 MJ | 0.19% |
| diet: total amount of ash | numerical | 1519.0±338.0 g | 0.19% |
| diet: total amount of crude fibre | numerical | 3619.0±581.0 g | 0.19% |
| diet: total amount of ether extracts | numerical | 601.0±158.0 g | 0.19% |
| diet: total amount of undegraded dietary protein | numerical | 598.0±226.0 g | 0.19% |
| diet: percentage of undegraded dietary protein | numerical | 0.0±0.0 % | 0.00% |
| diet: total ruminal nitrogen balance | numerical | 26.6±40.5 g | 0.19% |
| diet: content of acid detergent fiber | numerical | 0.0±0.0 g/kg dry mass | 0.38% |
| diet: content of acid detergent lignin | numerical | 0.0±0.0 g/kg dry mass | 0.38% |
| diet: content of metabolisable energy | numerical | 10.6±0.5 MJ/kg dry mass | 0.38% |
| diet: content of nitrogen-free extracts | numerical | 528.0±36.0 g/kg dry mass | 0.38% |
| diet: content of utilizable protein | numerical | 144.0±10.0 g/kg dry mass | 0.38% |
| diet: content of organic matter | numerical | 916.0±14.0 g/kg dry mass | 0.38% |
| diet: content of ash | numerical | 84.4±14.3 g/kg dry mass | 0.38% |
| diet: content of crude fibre | numerical | 203.0±34.0 g/kg dry mass | 0.38% |
| diet: content of crude fat | numerical | 32.8±3.9 g/kg dry mass | 0.38% |
| diet: content of crude protein | numerical | 152.0±20.0 g/kg dry mass | 0.38% |
| diet: percentage content of undegraded protein | numerical | 0.0±0.0 % | 0.38% |
| concentrate dry matter intake | numerical | 13.4±2.1 | 0.38% |
| dry matter | numerical | 9.01±0.4 kg | 58.09% |

**Table 8.** Features summarised in the feature category *feed*: For categorical and binary features, the frequency of each category is reported. For numerical features, mean values and standard deviations are reported. For all features, the percentage of missing values in the data set is reported.

| age | | | |
|---|---|---|---|
| **feature** | **type** | **values** | **% missing** |
| parity | numerical | 2.8±2.05 | 0.00% |
| age at first calving | numerical | 874.0±103.0 days | 0.09% |

**Table 9.** Features summarised in the feature category *age*: For categorical and binary features, the frequency of each category is reported. For numerical features, mean values and standard deviations are reported. For all features, the percentage of missing values in the data set is reported.

| lactation stage | | | |
|---|---|---|---|
| **feature** | **type** | **values** | **% missing** |
| in dry period during DHI (MLP) | categorical | True 86.0%, False 14.0% | 0.0% |
| days pregnant at DHI (MLP) | numerical | 102.0±101.0 days | 0.00% |
| days in milk at DHI (MLP) | numerical | 185.0±131.0 days | 3.35% |

**Table 10.** Features summarised in the feature category *lactation stage*: For categorical and binary features, the frequency of each category is reported. For numerical features, mean values and standard deviations are reported. For all features, the percentage of missing values in the data set is reported.

| breed | | | |
|---|---|---|---|
| **feature** | **type** | **values** | **% missing** |
| main breed | categorical | Fleckvieh 52.63%, Brown Swiss 26.82%, Holstein 18.82% | 2.00% |
| ratio of foreign genes | numerical | 6.94±14.01 % | 0.01% |
| ratio of Angler Rotvieh genes | numerical | 0.06±2.13 % | 0.00% |
| ratio of Blonde d'Aquitaine genes | numerical | 0.02±0.99 % | 0.00% |
| ration of original Braunvieh genes | numerical | 0.28±3.92 % | 0.00% |
| ratio of Braunvieh genes | numerical | 0.32±3.68 % | 0.00% |
| ratio of meat Fleckvieh genes | numerical | 0.01±0.22 % | 0.00% |
| ratio of Fleckvieh genes | numerical | 1.75±8.13 % | 0.00% |
| ratio of Holstein genes | numerical | 0.13±2.31 % | 0.00% |
| ratio of Jersey genes | numerical | 0.08±1.48 % | 0.00% |
| ratio of Montbeliarde genes | numerical | 1.05±5.55 % | 0.00% |
| ratio of Pinzgauer genes | numerical | 0.05±1.73 % | 0.00% |
| ratio of Piemonteser genes | numerical | 0.02±0.99 % | 0.00% |
| ratio of Holstein Rotbunte genes | numerical | 4.86±11.23 % | 0.00% |

**Table 11.** Features summarised in the feature category *breed*: For categorical and binary features, the frequency of each category is reported. For numerical features, mean values and standard deviations are reported. For all features, the percentage of missing values in the data set is reported.

| breeding values | | | |
|---|---|---|---|
| **feature** | **type** | **values** | **% missing** |
| TMI | numerical | 87.2±9.1 | 2.55% |
| milk index | numerical | 87.5±9.4 | 1.98% |
| beef index | numerical | 97.7±9.1 | 21.07% |
| fitness index | numerical | 98.0±8.2 | 20.73% |
| breeding value: milk yield | numerical | -487.1±463.2 | 1.98% |

**Table 12.** Features summarised in the feature category *breeding values*: For categorical and binary features, the frequency of each category is reported. For numerical features, mean values and standard deviations are reported. For all features, the percentage of missing values in the data set is reported.

**environment**

| **feature** | **type** | **values** | **% missing** |
|---|---|---|---|
| season | categorical | autumn 27.61%, winter 26.65%, spring 22.12%, summer 23.61% | 0.00% |
| altitude | numerical | 605.0±231.0 m | 0.00% |
| mean yearly relative humidity | numerical | 80.4±2.9 % | 0.00% |
| standard deviation of yearly relative humidity | numerical | 11.7±1.5 % | 0.00% |
| mean yearly precipitation | numerical | 2.92±0.75 mm | 0.00% |
| standard deviation of yearly precipitation | numerical | 6.28±1.18 mm | 0.00% |
| mean yearly temperature | numerical | 10.3±1.3 C° | 0.00% |
| standard deviation of yearly temperature | numerical | 6.61±0.27 C° | 0.00% |
| number of high wind days | numerical | 0.1±0.12 days | 0.00% |
| number of low temperature days | numerical | 0.16±0.07 days | 0.00% |

**Table 13.** Features summarised in the feature category *environment*: For categorical and binary features, the frequency of each category is reported. For numerical features, mean values and standard deviations are reported. For all features, the percentage of missing values in the data set is reported.

**lameness**

| rank | feature | category | % | OR | $p$-value |
|---|---|---|---|---|---|
| 1 | parity | age | 10.62 ± 1.87 | 2.25 [2.17; 2.33] | <0.001 |
| 2 | diagnosis source: culling reason or observation at calving* | diagnoses | 5.94 ± 1.06 | 0.31 [0.29; 0.33] | <0.001 |
| 3 | diagnosis source: score* | diagnoses | 3.56 ± 0.98 | 1.53 [1.48; 1.59] | <0.001 |
| 4 | mean yearly precipitation | environment | 2.45 ± 0.79 | 0.80 [0.77; 0.84] | <0.001 |
| 5 | body condition score | physique | 2.34 ± 0.80 | 0.71 [0.68; 0.73] | <0.001 |
| 6 | standard deviation of yearly temperature | environment | 2.33 ± 0.81 | 1.40 [1.34; 1.46] | <0.001 |
| 7 | litter in free-stall for lactating cows: other | housing | 1.59 ± 0.60 | 0.80 [0.72; 0.88] | <0.001 |
| 8 | forage type: sequentially fed forages | feed | 1.49 ± 0.52 | 0.65 [0.60; 0.71] | <0.001 |
| 9 | dry matter | feed | 1.41 ± 0.51 | 0.94 [0.88; 1.00] | 0.034 |
| 10 | litter in free-stall for lactating cows: chopped straw | housing | 1.33 ± 0.59 | 1.54 [1.41; 1.69] | <0.001 |
| 11 | mean yearly temperature | environment | 1.21 ± 0.54 | 1.23 [1.18; 1.29] | <0.001 |
| 12 | automated milking switch-off | husbandry | 1.15 ± 0.44 | 0.70 [0.63; 0.77] | <0.001 |
| 13 | diet: content of organic matter | feed | 1.14 ± 0.46 | 0.90 [0.87; 0.94] | <0.001 |
| 14 | mean yearly relative humidity | environment | 1.00 ± 0.57 | 1.09 [1.05; 1.14] | <0.001 |
| 15 | number of low temperature days | environment | 0.98 ± 0.70 | 0.89 [0.86; 0.93] | <0.001 |
| 16 | standard deviation of yearly precipitation | environment | 0.96 ± 0.45 | 0.99 [0.95; 1.03] | 0.567 |
| 17 | TMI | breeding values | 0.96 ± 0.95 | 0.78 [0.74; 0.81] | <0.001 |
| 18 | claw trimming frequency: three times per year | husbandry | 0.93 ± 0.27 | 1.18 [1.06; 1.32] | 0.003 |
| 19 | type of milking stalls: pipe milking | housing | 0.91 ± 0.43 | 0.47 [0.39; 0.56] | <0.001 |
| 20 | milking vacuum | husbandry | 0.90 ± 0.40 | 0.87 [0.84; 0.91] | <0.001 |
| 21 | annual herd milk yield average* | husbandry | 0.88 ± 0.52 | 0.88 [0.84; 0.92] | <0.001 |
| 22 | dietary proportion of hay | feed | 0.87 ± 0.49 | 0.95 [0.91; 0.99] | 0.017 |
| 23 | floor in open air area for young stock: no open-air areas | housing | 0.86 ± 0.47 | 1.88 [1.72; 2.06] | <0.001 |
| 24 | number of high wind days | environment | 0.85 ± 0.63 | 1.18 [1.13; 1.22] | <0.001 |
| 25 | standard deviation of yearly relative humidity | environment | 0.84 ± 0.57 | 1.07 [1.03; 1.11] | <0.001 |
| 26 | used forage types in diet: field forage silage, grass silage, hay, corn silage | feed | 0.81 ± 0.33 | 1.17 [1.06; 1.30] | 0.002 |
| 27 | diet: content of crude protein | feed | 0.81 ± 0.46 | 1.16 [1.11; 1.21] | <0.001 |
| 28 | free-stall system for lactating cows: deep bed cubicle | housing | 0.78 ± 0.40 | 0.39 [0.36; 0.43] | <0.001 |
| 29 | free-stall system for dry cows: deep litter | housing | 0.78 ± 0.21 | 0.56 [0.48; 0.64] | <0.001 |
| 30 | manure removal in free-stall for dry cows: slits | housing | 0.78 ± 0.40 | 2.15 [1.97; 2.35] | <0.001 |
| 31 | diet: total amount of ether extracts | feed | 0.76 ± 0.43 | 1.10 [1.05; 1.15] | <0.001 |
| 32 | floor in open-air area for lactating cows: no open-air areas | housing | 0.76 ± 0.41 | 1.85 [1.71; 2.01] | <0.001 |
| 33 | herd size | husbandry | 0.75 ± 0.49 | 1.16 [1.11; 1.21] | <0.001 |
| 34 | claw trimming frequency: only for lame animals | husbandry | 0.75 ± 0.23 | 0.86 [0.72; 1.02] | 0.085 |
| 35 | litter in free-stall for dry cows: chopped straw | housing | 0.74 ± 0.45 | 1.63 [1.49; 1.79] | <0.001 |
| 36 | lactating cows on alpine pasture | husbandry | 0.74 ± 0.61 | 0.19 [0.15; 0.24] | <0.001 |
| 37 | pasture of lactating cows | husbandry | 0.73 ± 0.48 | 0.47 [0.43; 0.52] | <0.001 |
| 38 | chest girth | physique | 0.71 ± 0.73 | 1.26 [1.20; 1.32] | <0.001 |
| 39 | floor in walkway of free-stall for young stock: concrete slits | housing | 0.69 ± 0.32 | 2.29 [2.10; 2.50] | <0.001 |
| 40 | manure removal in free-stall for young stock: other | housing | 0.66 ± 0.38 | 0.53 [0.49; 0.58] | <0.001 |
| 41 | walkway floor in free-stall for dry cows: concrete slits | housing | 0.65 ± 0.24 | 1.55 [1.41; 1.70] | <0.001 |
| 42 | farm organically managed | husbandry | 0.65 ± 0.29 | 0.61 [0.54; 0.68] | <0.001 |
| 43 | dietary proportion of grass silage | feed | 0.63 ± 0.48 | 0.96 [0.92; 1.00] | 0.036 |
| 44 | diet: content of utilizable protein | feed | 0.62 ± 0.37 | 1.21 [1.15; 1.27] | <0.001 |
| 45 | lying mats in free-stall for dry cows | housing | 0.62 ± 0.35 | 2.40 [2.17; 2.66] | <0.001 |
| 46 | muscularity score | physique | 0.61 ± 0.62 | 0.62 [0.60; 0.65] | <0.001 |
| 47 | floor in open-air area for dry cows: no open-air areas | housing | 0.60 ± 0.35 | 1.36 [1.25; 1.48] | <0.001 |
| 48 | walkway floor in free-stall for lactating cows: rubber mats | housing | 0.60 ± 0.31 | 1.12 [1.00; 1.24] | 0.043 |
| 49 | diet: content of crude fibre | feed | 0.60 ± 0.37 | 0.78 [0.74; 0.82] | <0.001 |
| 50 | claw trimming done by farmer | husbandry | 0.59 ± 0.30 | 0.79 [0.72; 0.86] | <0.001 |

**Table 14.** Feature importances and odds ratios including $p$-values for the 50 most important features for lameness. Features indicated with a star are treated as covariates in the logistic regression and their odds ratios have been calculated separately. Odds ratios $\geq 1.1$ and $\leq 0.9$ are highlighted with yellow and green cell backgrounds respectively.

**anoestrus**

| rank | feature | category | % | OR | $p$-value |
|---|---|---|---|---|---|
| 1 | diagnosis source: performance recording organisation (LKV)* | diagnoses | 5.60 ± 0.94 | 2.78 [2.63; 2.93] | <0.001 |
| 2 | diet: total amount of crude fibre | feed | 3.45 ± 0.77 | 0.85 [0.77; 0.93] | <0.001 |
| 3 | barn design: outdoor climate house open front | housing | 3.41 ± 0.99 | 3.20 [2.68; 3.82] | <0.001 |
| 4 | lactating cows, time on pasture | husbandry | 3.25 ± 0.67 | 0.81 [0.62; 1.07] | 0.135 |
| 5 | diet: content of nitrogen-free extracts | feed | 2.72 ± 0.79 | 1.96 [1.78; 2.17] | <0.001 |
| 6 | used forage types in diet: field forage silage, grass silage, hay, corn silage, pasture | feed | 2.42 ± 0.74 | 1.72 [1.40; 2.13] | <0.001 |
| 7 | floor in open-air area for lactating cows: no open-air areas | housing | 2.41 ± 1.19 | 2.95 [2.48; 3.52] | <0.001 |
| 8 | diagnosis source: culling reason or observation at calving* | diagnoses | 2.17 ± 1.82 | 0.11 [0.09; 0.14] | <0.001 |
| 9 | breeding value: milk yield | breeding values | 2.17 ± 1.22 | 1.34 [1.23; 1.47] | <0.001 |
| 10 | diet: content of metabolisable energy | feed | 2.15 ± 1.21 | 1.66 [1.46; 1.88] | <0.001 |
| 11 | test-day milk yield | milk | 2.08 ± 1.16 | 1.52 [1.37; 1.70] | <0.001 |
| 12 | manure removal in free-stall for young stock: other | housing | 1.98 ± 1.27 | 1.37 [1.15; 1.63] | <0.001 |
| 13 | fitness index | breeding values | 1.90 ± 0.97 | 0.76 [0.69; 0.83] | <0.001 |
| 14 | number of high wind days | environment | 1.76 ± 1.06 | 1.29 [1.19; 1.40] | <0.001 |
| 15 | annual herd milk yield average* | husbandry | 1.75 ± 1.11 | 1.65 [1.50; 1.80] | <0.001 |
| 16 | diet: content of ash | feed | 1.70 ± 1.19 | 0.65 [0.58; 0.72] | <0.001 |
| 17 | mean yearly precipitation | environment | 1.68 ± 1.28 | 0.50 [0.44; 0.57] | <0.001 |
| 18 | diet: content of utilizable protein | feed | 1.63 ± 1.14 | 1.45 [1.30; 1.62] | <0.001 |
| 19 | forage type: grass plus corn and grass products plus corn | feed | 1.50 ± 0.55 | 2.39 [1.85; 3.07] | <0.001 |
| 20 | floor in open-air area for dry cows: no open-air areas | housing | 1.38 ± 1.11 | 3.83 [3.08; 4.77] | <0.001 |
| 21 | body condition score | physique | 1.36 ± 0.28 | 0.83 [0.76; 0.90] | <0.001 |
| 22 | used forage types in diet: grass silage, hay | feed | 1.34 ± 0.32 | 0.46 [0.31; 0.67] | <0.001 |
| 23 | herd size | husbandry | 1.33 ± 0.87 | 1.37 [1.26; 1.48] | <0.001 |
| 24 | test-day energy corrected milk daily yield | milk | 1.32 ± 0.34 | 0.97 [0.90; 1.04] | 0.344 |
| 25 | lactating cows are kept in a tie stall facility | husbandry | 1.30 ± 0.40 | 0.85 [0.58; 1.24] | 0.387 |
| 26 | manure removal in free-stall for young stock: scraper | housing | 1.24 ± 0.46 | 0.53 [0.41; 0.67] | <0.001 |
| 27 | free-stall system for young stock: deep litter | housing | 1.24 ± 0.46 | 2.48 [2.07; 2.99] | <0.001 |
| 28 | type of milking stalls: milking robot | housing | 1.17 ± 0.47 | 0.57 [0.45; 0.71] | <0.001 |
| 29 | floor in walkway of free-stall for young stock: solid concrete with slits | housing | 1.16 ± 0.54 | 1.79 [1.34; 2.40] | <0.001 |
| 30 | diet: total amount of undegraded dietary protein | feed | 1.12 ± 0.91 | 1.74 [1.58; 1.92] | <0.001 |
| 31 | floor in open air area for young stock: no open-air areas | housing | 1.11 ± 0.43 | 1.84 [1.52; 2.24] | <0.001 |
| 32 | floor in open-air areas for lactating cows: solid concrete | housing | 1.11 ± 1.02 | 0.22 [0.18; 0.27] | <0.001 |
| 33 | claw trimming frequency: once per year | husbandry | 1.10 ± 0.59 | 1.29 [1.02; 1.63] | 0.033 |
| 34 | parity | age | 1.09 ± 0.76 | 1.45 [1.34; 1.58] | <0.001 |
| 35 | free-stall system for young stock: sloped floor | housing | 1.05 ± 0.57 | 0.60 [0.37; 0.96] | 0.034 |
| 36 | free-stall system for young stock: high bed cubicle | housing | 1.02 ± 0.47 | 1.02 [0.86; 1.22] | 0.822 |
| 37 | mean yearly relative humidity | environment | 0.93 ± 0.68 | 1.05 [0.96; 1.16] | 0.260 |
| 38 | days in milk at DHI (MLP)* | lactation stage | 0.90 ± 0.74 | 0.61 [0.56; 0.68] | <0.001 |
| 39 | dietary proportion of grass silage | feed | 0.88 ± 0.62 | 0.61 [0.56; 0.67] | <0.001 |
| 40 | claw trimming frequency: twice per year | husbandry | 0.87 ± 0.68 | 0.57 [0.48; 0.68] | <0.001 |
| 41 | dietary proportion of concentrates | feed | 0.84 ± 0.60 | 1.42 [1.27; 1.60] | <0.001 |
| 42 | litter in free-stall for dry cows: chopped straw | housing | 0.82 ± 0.68 | 1.03 [0.86; 1.23] | 0.727 |
| 43 | diagnosis source: score* | diagnoses | 0.82 ± 1.13 | 0.42 [0.39; 0.46] | <0.001 |
| 44 | forage type: partial mixed ration | feed | 0.80 ± 0.69 | 0.56 [0.46; 0.70] | <0.001 |
| 45 | problematic feed quality | feed | 0.79 ± 0.70 | 1.52 [1.19; 1.93] | <0.001 |
| 46 | floor in open-air area for young stock: unpaved | housing | 0.78 ± 0.35 | 0.31 [0.19; 0.49] | <0.001 |
| 47 | walkway floor in free-stall for dry cows: other | housing | 0.75 ± 0.45 | 0.90 [0.73; 1.10] | 0.284 |
| 48 | test-day protein yield percentage | milk | 0.74 ± 0.70 | 0.71 [0.64; 0.80] | <0.001 |
| 49 | diet: total amount of ether extracts | feed | 0.74 ± 0.65 | 1.44 [1.32; 1.57] | <0.001 |
| 50 | season: autumn* | environment | 0.72 ± 0.48 | 2.04 [1.76; 2.36] | <0.001 |

**Table 15.** Feature importances and odds ratios including $p$-values for the 50 most important features for anoestrus. Features indicated with a star are treated as covariates in the logistic regression and their odds ratios have been calculated separately. Odds ratios $\geq 1.1$ and $\leq 0.9$ are highlighted with yellow and green cell backgrounds respectively.

**ketosis**

| rank | feature | category | % | OR | $p$-value |
|---|---|---|---|---|---|
| 1 | days in milk at DHI (MLP)* | lactation stage | $3.77 \pm 0.94$ | 1.23 [1.14; 1.33] | <0.001 |
| 2 | test-day energy corrected milk daily yield | milk | $2.31 \pm 1.39$ | 1.01 [0.89; 1.16] | 0.843 |
| 3 | standard deviation of yearly temperature | environment | $2.28 \pm 0.84$ | 1.89 [1.71; 2.09] | <0.001 |
| 4 | in dry period during DHI (MLP) | lactation stage | $2.20 \pm 0.62$ | excuded | |
| 5 | herd size | husbandry | $1.97 \pm 0.73$ | 1.37 [1.26; 1.48] | <0.001 |
| 6 | litter in free-stall for young stock: chopped straw | housing | $1.76 \pm 0.74$ | 2.56 [2.02; 3.25] | <0.001 |
| 7 | mean yearly temperature | environment | $1.65 \pm 0.77$ | 1.23 [1.11; 1.35] | <0.001 |
| 8 | diagnosis source: culling reason or observation at calving* | diagnoses | $1.51 \pm 1.07$ | 0.38 [0.33; 0.43] | <0.001 |
| 9 | dietary proportion of grass silage | feed | $1.50 \pm 0.68$ | 0.90 [0.83; 0.97] | 0.008 |
| 10 | problematic feed quality | feed | $1.50 \pm 0.63$ | 1.26 [1.00; 1.60] | 0.054 |
| 11 | body weight | physique | $1.49 \pm 0.73$ | 2.38 [2.14; 2.64] | <0.001 |
| 12 | free-stall system for young stock: other | housing | $1.45 \pm 0.48$ | 1.64 [1.32; 2.03] | <0.001 |
| 13 | diagnosis source: veterinarian | diagnoses | $1.44 \pm 0.74$ | 1.53 [1.43; 1.64] | <0.001 |
| 14 | number of low temperature days | environment | $1.44 \pm 0.71$ | 0.95 [0.87; 1.03] | 0.204 |
| 15 | muscularity score | physique | $1.40 \pm 0.62$ | 1.02 [0.93; 1.12] | 0.713 |
| 16 | dietary proportion of concentrates | feed | $1.36 \pm 0.69$ | 0.57 [0.51; 0.63] | <0.001 |
| 17 | chest girth | physique | $1.35 \pm 0.88$ | 1.96 [1.79; 2.14] | <0.001 |
| 18 | altitude | environment | $1.34 \pm 0.66$ | 0.74 [0.67; 0.81] | <0.001 |
| 19 | diet: content of ash | feed | $1.31 \pm 0.77$ | 0.92 [0.84; 1.00] | 0.053 |
| 20 | diet: content of crude fibre | feed | $1.30 \pm 0.78$ | 1.63 [1.50; 1.77] | <0.001 |
| 21 | used forage types in diet: field forage silage, grass silage, hay, corn silage | feed | $1.29 \pm 0.51$ | 2.24 [1.86; 2.70] | <0.001 |
| 22 | mean yearly precipitation | environment | $1.26 \pm 0.82$ | 0.46 [0.41; 0.52] | <0.001 |
| 23 | standard deviation of yearly relative humidity | environment | $1.26 \pm 0.85$ | 0.80 [0.73; 0.87] | <0.001 |
| 24 | diet: total amount of ash | feed | $1.23 \pm 0.81$ | 0.49 [0.44; 0.54] | <0.001 |
| 25 | diet: total amount of ether extracts | feed | $1.22 \pm 0.66$ | 0.45 [0.41; 0.51] | <0.001 |
| 26 | diet: content of organic matter | feed | $1.21 \pm 0.55$ | 1.08 [0.99; 1.18] | 0.090 |
| 27 | mean yearly relative humidity | environment | $1.20 \pm 0.64$ | 1.39 [1.26; 1.54] | <0.001 |
| 28 | number of milking places | husbandry | $1.15 \pm 0.63$ | 1.07 [0.99; 1.15] | 0.078 |
| 29 | test-day milk yield | milk | $1.14 \pm 1.05$ | 0.80 [0.67; 0.97] | 0.022 |
| 30 | floor in walkway of free-stall for young stock walkway: other | housing | $1.14 \pm 0.48$ | 0.89 [0.70; 1.13] | 0.338 |
| 31 | TMI | breeding values | $1.07 \pm 0.67$ | 0.87 [0.79; 0.96] | 0.004 |
| 32 | milk index | breeding values | $1.04 \pm 0.62$ | 1.03 [0.94; 1.13] | 0.520 |
| 33 | diagnosis source: score* | diagnoses | $0.99 \pm 0.64$ | 0.83 [0.77; 0.90] | <0.001 |
| 34 | test-day protein yield percentage | milk | $0.97 \pm 0.83$ | 1.43 [1.25; 1.65] | <0.001 |
| 35 | diet: content of nitrogen-free extracts | feed | $0.96 \pm 0.67$ | 0.85 [0.78; 0.93] | <0.001 |
| 36 | manure removal: slurry with solid flooring | housing | $0.96 \pm 0.64$ | 1.27 [1.08; 1.50] | 0.004 |
| 37 | walkway floor in free-stall for dry cows: other | housing | $0.92 \pm 0.54$ | 0.62 [0.49; 0.79] | <0.001 |
| 38 | provision of supplementary concentrate: other | feed | $0.92 \pm 0.26$ | 2.54 [2.07; 3.10] | <0.001 |
| 39 | waist circumference | physique | $0.90 \pm 0.62$ | 1.58 [1.43; 1.75] | <0.001 |
| 40 | ration type: total mixed ration | feed | $0.89 \pm 0.19$ | 3.78 [2.96; 4.83] | <0.001 |
| 41 | diet: total amount of undegraded dietary protein | feed | $0.88 \pm 0.72$ | 0.48 [0.43; 0.54] | <0.001 |
| 42 | forage type: grass and grass products plus corn only | feed | $0.86 \pm 0.23$ | 1.07 [0.77; 1.48] | 0.698 |
| 43 | test-day urea content | milk | $0.86 \pm 0.62$ | 0.89 [0.78; 1.02] | 0.091 |
| 44 | annual herd milk yield average* | husbandry | $0.84 \pm 0.65$ | 1.08 [0.98; 1.19] | 0.115 |
| 45 | forage type: year-round silage (with corn silage) | feed | $0.83 \pm 0.29$ | 1.43 [1.15; 1.77] | 0.001 |
| 46 | test-day somatic cell count | milk | $0.83 \pm 0.93$ | 1.63 [1.46; 1.82] | <0.001 |
| 47 | lying mats in free-stall for young stock | housing | $0.81 \pm 0.48$ | 1.33 [1.08; 1.63] | 0.007 |
| 48 | diet: content of metabolisable energy | feed | $0.80 \pm 0.68$ | 0.64 [0.59; 0.70] | <0.001 |
| 49 | diet: total ruminal nitrogen balance | feed | $0.79 \pm 0.68$ | 0.74 [0.68; 0.81] | <0.001 |
| 50 | litter in free-stall for lactating cows: chopped straw | housing | $0.78 \pm 0.67$ | 1.80 [1.49; 2.17] | <0.001 |

**Table 16.** Feature importances and odds ratios including $p$-values for the 50 most important features for ketosis. Features indicated with a star are treated as covariates in the logistic regression and their odds ratios have been calculated separately. Odds ratios $\geq 1.1$ and $\leq 0.9$ are highlighted with yellow and green cell backgrounds respectively.

**periparturient hypocalcemia**

| rank | feature | category | % | OR | *p*-value |
|---|---|---|---|---|---|
| 1 | parity | age | 19.49 ± 8.13 | 2.12 [1.98; 2.27] | <0.001 |
| 2 | days in milk at DHI (MLP)* | lactation stage | 9.02 ± 4.64 | 2.20 [2.03; 2.37] | <0.001 |
| 3 | days pregnant at DHI (MLP) | lactation stage | 8.66 ± 3.62 | excuded | |
| 4 | in dry period during DHI (MLP) | lactation stage | 8.18 ± 3.53 | excuded | |
| 5 | test-day protein yield percentage | milk | 4.24 ± 3.62 | 1.26 [1.09; 1.45] | 0.002 |
| 6 | used forage types in diet: field forage silage, grass silage, hay, corn silage | feed | 3.73 ± 1.81 | 0.93 [0.74; 1.16] | 0.505 |
| 7 | test-day somatic cell count | milk | 3.61 ± 3.48 | 1.07 [0.94; 1.22] | 0.320 |
| 8 | test-day lactose content | milk | 2.61 ± 3.07 | 0.93 [0.82; 1.05] | 0.231 |
| 9 | waist circumference | physique | 1.85 ± 2.62 | 1.79 [1.62; 1.97] | <0.001 |
| 10 | diet: total amount of crude fibre | feed | 1.82 ± 1.91 | 0.70 [0.64; 0.75] | <0.001 |
| 11 | test-day fat yield percentage | milk | 1.75 ± 2.34 | 1.15 [1.01; 1.30] | 0.031 |
| 12 | barn design: other | housing | 1.52 ± 2.26 | excuded | |
| 13 | mean yearly relative humidity | environment | 1.44 ± 2.13 | 1.17 [1.06; 1.30] | 0.002 |
| 14 | body condition score | physique | 1.29 ± 0.94 | 1.24 [1.14; 1.36] | <0.001 |
| 15 | body weight | physique | 1.09 ± 1.83 | 1.70 [1.53; 1.88] | <0.001 |
| 16 | diet: total amount of undegraded dietary protein | feed | 1.02 ± 0.97 | 0.63 [0.56; 0.70] | <0.001 |
| 17 | test-day milk yield | milk | 0.94 ± 1.36 | 1.24 [1.04; 1.49] | 0.018 |
| 18 | breeding value: milk yield | breeding values | 0.93 ± 0.92 | 1.16 [1.06; 1.28] | 0.002 |
| 19 | pasture duration of dry cows | husbandry | 0.89 ± 0.73 | 1.02 [0.87; 1.20] | 0.794 |
| 20 | litter in free-stall for lactating cows: chopped straw | housing | 0.82 ± 0.63 | 1.26 [1.05; 1.51] | 0.012 |
| 21 | walkway floor in free-stall for dry cows: solid concrete | housing | 0.82 ± 0.53 | 1.12 [0.94; 1.35] | 0.211 |
| 22 | litter in free-stall for dry cows: long straw | housing | 0.81 ± 0.63 | 0.87 [0.70; 1.08] | 0.210 |
| 23 | ration type: feedstuffs sequentially fed | feed | 0.80 ± 0.61 | 0.97 [0.82; 1.15] | 0.763 |
| 24 | number of milking places | husbandry | 0.78 ± 1.18 | 1.13 [1.04; 1.23] | 0.004 |
| 25 | diet: content of nitrogen-free extracts | feed | 0.70 ± 0.88 | 0.95 [0.88; 1.03] | 0.250 |
| 26 | chest girth | physique | 0.69 ± 1.50 | 1.27 [1.16; 1.40] | <0.001 |
| 27 | diet: content of crude protein | feed | 0.68 ± 0.92 | 0.76 [0.70; 0.82] | <0.001 |
| 28 | beef index | breeding values | 0.63 ± 0.77 | 0.91 [0.84; 0.99] | 0.025 |
| 29 | test-day urea content | milk | 0.56 ± 1.24 | 0.95 [0.83; 1.08] | 0.411 |
| 30 | dry cows on alpine pasture | husbandry | 0.55 ± 1.27 | 0.63 [0.44; 0.90] | 0.011 |
| 31 | type of milking stalls: milking robot | housing | 0.53 ± 0.43 | 0.63 [0.49; 0.83] | <0.001 |
| 32 | farm organically managed | husbandry | 0.49 ± 0.44 | 0.70 [0.55; 0.89] | 0.004 |
| 33 | standard deviation of yearly temperature | environment | 0.47 ± 1.01 | 1.48 [1.34; 1.63] | <0.001 |
| 34 | ration type: partial mixed ration | feed | 0.43 ± 0.55 | 0.92 [0.78; 1.09] | 0.350 |
| 35 | barn design: free-stall barn | housing | 0.43 ± 0.44 | 1.01 [0.86; 1.19] | 0.903 |
| 36 | provision of supplementary concentrate: electronic feeder | feed | 0.42 ± 0.51 | 0.70 [0.59; 0.84] | <0.001 |
| 37 | forage type: partial mixed ration | feed | 0.42 ± 0.55 | 0.79 [0.65; 0.96] | 0.017 |
| 38 | dietary proportion of clover | feed | 0.39 ± 0.57 | 0.66 [0.50; 0.87] | 0.003 |
| 39 | free-stall system for dry cows: high bed cubicle | housing | 0.37 ± 0.51 | 1.44 [1.11; 1.88] | 0.007 |
| 40 | litter in free-stall for dry cows: chopped straw | housing | 0.36 ± 0.43 | 1.42 [1.18; 1.71] | <0.001 |
| 41 | milking take off: present | husbandry | 0.34 ± 0.42 | 1.01 [0.86; 1.20] | 0.882 |
| 42 | litter in free-stall for young stock: chopped straw | housing | 0.33 ± 0.63 | 1.94 [1.52; 2.48] | <0.001 |
| 43 | free-stall system for young stock: other | housing | 0.33 ± 0.51 | 1.40 [1.13; 1.74] | 0.002 |
| 44 | walkway floor in free-stall for lactating cows: rubber mats | housing | 0.33 ± 0.48 | 0.82 [0.64; 1.06] | 0.131 |
| 45 | manure removal in free-stall for lactating cows: slits | housing | 0.33 ± 0.42 | 1.04 [0.86; 1.25] | 0.708 |
| 46 | manure removal in free-stall for dry cows: slits | housing | 0.32 ± 0.52 | 1.54 [1.27; 1.87] | <0.001 |
| 47 | manure removal in free-stall for dry cows: scraper | housing | 0.31 ± 0.50 | 0.86 [0.72; 1.03] | 0.109 |
| 48 | forage type: mixed ration with concentrates | feed | 0.30 ± 0.42 | 0.99 [0.83; 1.19] | 0.953 |
| 49 | manure removal: slurry with perforated flooring | housing | 0.30 ± 0.49 | 1.05 [0.88; 1.25] | 0.569 |
| 50 | age at first calving | age | 0.29 ± 0.59 | 0.97 [0.88; 1.06] | 0.469 |

**Table 17.** Feature importances and odds ratios including *p*-values for the 50 most important features for periparturient hypocalcemia. Features indicated with a star are treated as covariates in the logistic regression and their odds ratios have been calculated separately. Odds ratios $\geq 1.1$ and $\leq 0.9$ are highlighted with yellow and green cell backgrounds respectively.

**metritis**

| rank | feature | category | % | OR | *p*-value |
|---|---|---|---|---|---|
| 1 | diagnosis source: performance recording organisation (LKV)* | diagnoses | 19.46 ± 5.87 | 2.58 [2.40; 2.77] | <0.001 |
| 2 | ration type: feedstuffs sequentially fed | feed | 6.50 ± 3.87 | 0.35 [0.26; 0.46] | <0.001 |
| 3 | diagnosis source: score* | diagnoses | 4.73 ± 5.67 | 0.44 [0.39; 0.50] | <0.001 |
| 4 | diagnosis source: culling reason or observation at calving* | diagnoses | 4.15 ± 5.07 | 0.15 [0.12; 0.20] | <0.001 |
| 5 | standard deviation of yearly temperature | environment | 3.66 ± 4.78 | 2.94 [2.45; 3.53] | <0.001 |
| 6 | body weight | physique | 2.92 ± 2.69 | 1.42 [1.22; 1.66] | <0.001 |
| 7 | test-day milk yield | milk | 2.81 ± 1.40 | 0.99 [0.83; 1.19] | 0.933 |
| 8 | parity | age | 2.22 ± 1.76 | 1.72 [1.55; 1.91] | <0.001 |
| 9 | test-day energy corrected milk daily yield | milk | 1.81 ± 2.11 | 1.03 [0.91; 1.15] | 0.655 |
| 10 | number of high wind days | environment | 1.65 ± 1.65 | 1.10 [0.98; 1.24] | 0.112 |
| 11 | breeding value: milk yield | breeding values | 1.64 ± 1.18 | 1.13 [1.00; 1.29] | 0.056 |
| 12 | dietary proportion of clover | feed | 1.58 ± 0.73 | 1.40 [0.97; 2.02] | 0.072 |
| 13 | days pregnant at DHI (MLP) | lactation stage | 1.52 ± 0.93 | excuded | |
| 14 | manure removal in free-stall for dry cows: other | housing | 1.47 ± 1.62 | 1.41 [1.10; 1.80] | 0.007 |
| 15 | ration type: partial mixed ration | feed | 1.47 ± 1.12 | 1.72 [1.35; 2.19] | <0.001 |
| 16 | floor in open-air area for dry cows: no open-air areas | housing | 1.30 ± 1.74 | 3.12 [2.33; 4.18] | <0.001 |
| 17 | test-day protein yield percentage | milk | 1.22 ± 0.73 | 0.88 [0.74; 1.05] | 0.149 |
| 18 | waist circumference | physique | 1.18 ± 1.25 | 1.17 [1.01; 1.36] | 0.033 |
| 19 | season: summer* | environment | 1.18 ± 0.77 | 0.51 [0.37; 0.70] | <0.001 |
| 20 | season: autumn* | environment | 1.17 ± 1.23 | 2.38 [1.90; 2.98] | <0.001 |
| 21 | season: spring* | environment | 1.17 ± 0.76 | 0.71 [0.53; 0.94] | 0.018 |
| 22 | test-day urea content | milk | 1.13 ± 0.82 | 0.85 [0.73; 0.99] | 0.032 |
| 23 | manure removal in free-stall for dry cows: scraper | housing | 1.13 ± 1.68 | 0.49 [0.38; 0.65] | <0.001 |
| 24 | litter in free-stall for lactating cows: chopped straw | housing | 1.04 ± 2.04 | 2.42 [1.81; 3.23] | <0.001 |
| 25 | TMI | breeding values | 1.01 ± 0.82 | 0.85 [0.75; 0.98] | 0.023 |
| 26 | chest girth | physique | 1.00 ± 1.64 | 1.38 [1.21; 1.58] | <0.001 |
| 27 | dietary proportion of straw | feed | 0.88 ± 0.97 | 1.40 [1.10; 1.79] | 0.007 |
| 28 | walkway floor in free-stall for lactating cows: solid concrete with slits | housing | 0.88 ± 1.80 | 0.32 [0.18; 0.58] | <0.001 |
| 29 | farm organically managed | husbandry | 0.82 ± 1.80 | 0.33 [0.20; 0.55] | <0.001 |
| 30 | walkway floor in free-stall for lactating cows: rubberised slits | housing | 0.82 ± 1.67 | excuded | |
| 31 | free-stall system for dry cows: other | housing | 0.81 ± 1.81 | 0.48 [0.31; 0.76] | 0.002 |
| 32 | test-day somatic cell count | milk | 0.77 ± 0.82 | 1.09 [0.95; 1.25] | 0.201 |
| 33 | dietary proportion of concentrates | feed | 0.77 ± 0.81 | 0.84 [0.73; 0.98] | 0.025 |
| 34 | diet: total amount of crude fibre | feed | 0.75 ± 0.82 | 0.74 [0.65; 0.84] | <0.001 |
| 35 | litter in free-stall for lactating cows: long straw | housing | 0.75 ± 1.63 | | |
| 36 | diet: content of crude protein | feed | 0.74 ± 0.86 | 0.84 [0.74; 0.96] | 0.011 |
| 37 | lactating cows, time on pasture | husbandry | 0.73 ± 1.71 | 0.89 [0.63; 1.25] | 0.489 |
| 38 | diet: content of utilizable protein | feed | 0.71 ± 0.98 | 0.83 [0.72; 0.96] | 0.012 |
| 39 | diet: content of nitrogen-free extracts | feed | 0.68 ± 1.01 | 1.20 [1.05; 1.38] | 0.007 |
| 40 | diet: content of crude fat | feed | 0.66 ± 0.92 | 0.99 [0.88; 1.11] | 0.833 |
| 41 | forage type: grass and grass products only | feed | 0.66 ± 1.40 | excuded | |
| 42 | number of low temperature days | environment | 0.61 ± 1.14 | 0.67 [0.57; 0.78] | <0.001 |
| 43 | claw trimming frequency: twice per year | husbandry | 0.60 ± 0.79 | 0.57 [0.45; 0.73] | <0.001 |
| 44 | problematic feed quality | feed | 0.59 ± 0.81 | | |
| 45 | mean yearly precipitation | environment | 0.56 ± 1.60 | 0.59 [0.49; 0.70] | <0.001 |
| 46 | ratio of foreign genes | breed | 0.54 ± 0.85 | 0.93 [0.82; 1.04] | 0.193 |
| 47 | diet: total amount of ash | feed | 0.54 ± 0.82 | 0.75 [0.66; 0.86] | <0.001 |
| 48 | feeding group: lactating and dry cows on same forage (mixture) | feed | 0.52 ± 1.08 | excuded | |
| 49 | lactating cows on alpine pasture | husbandry | 0.52 ± 1.33 | excuded | |
| 50 | diet: total amount of undegraded dietary protein | feed | 0.51 ± 0.89 | 0.85 [0.74; 0.98] | 0.023 |

**Table 18.** Feature importances and odds ratios including *p*-values for the 50 most important features for metritis. Features indicated with a star are treated as covariates in the logistic regression and their odds ratios have been calculated separately. Odds ratios $\geq 1.1$ and $\leq 0.9$ are highlighted with yellow and green cell backgrounds respectively.
